# 3D imaging with superior resolution using Atacama Clear

**DOI:** 10.1101/2024.01.22.576689

**Authors:** Lauretta A. Lacko, Tansol Choi, Neranjan de Silva, Ying Liu, Edgar A. Jamies, Todd Evans, Romulo Hurtado

## Abstract

3-dimensional (3D) imaging is a powerful tool for interrogation of intact tissues, but can suffer from poor resolution due to impediments such as high tissue autofluorescence that remains a significant challenge in imaging cleared samples, including human clinical specimens. We developed Atacama Clear (ATC), a 3D imaging technology that increases signal-to-noise ratios (SNRs) while simultaneously augmenting the capacity of tissue to be cleared. ATC exhibited SNRs that are up to 200% of widely used 3D imaging methods, potentiated all tested optical clearing solutions by up to 600%, decreased the time of optical clearing by up to a factor of 8, and enabled detection of poorly recognized antigens with a remarkable 4-fold increase in signal detection while using up to 10-fold lower antibody concentrations. Strikingly, ATC produced up to a 5x increase in transgenic fluorescent reporter protein signal detection, which is instead often diminished with currently used 3D imaging methods. This increased imaging efficacy enabled multiplex interrogation of tough fibrous tissue and specimens that naturally exhibit high levels of background noise, including the heart, kidney, and human biopsies. Indeed, ATC facilitated the use of AI based auto-segmentation with simple low tech stereo fluorescence microscopy, visualization of previously undocumented adjacent nephron segments that exhibit notoriously high autofluorescence, elements of the cardiac conduction system, and distinct human glomerular tissue layers, with cellular resolution. Taken together, these studies establish ATC as a platform for complex 3D imaging studies of basic and clinical specimens with superior resolution.

## INTRODUCTION

Understanding the global tissue architecture of organs provides key insight into normal and pathological organ physiology. However, visualizing the 3-dimensional (3D) organization of tissue is limited by its natural opacity that causes light scattering and prevents deep penetration of light. Traditionally, these challenges have been addressed by assaying serial thin histological sections, and then attempting to derive 3D information. This approach, however, is laborious and time consuming, and can result in unsatisfactory reconstruction in the case of damaged or lost sections. To this end, our understanding of systems-level organ structure has been greatly advanced by optical clearing methods that enable direct 3D imaging through intact organs^1,2^.

In general, optical clearing methodologies consist of a lipid removal step that makes the whole tissue accessible, and a step immersing the tissue in refractive index matching solutions that allow light to travel through the specimen without light scatter. These protocols can be broadly categorized into either solvent-based or aqueous-based protocols^1,2^. Aqueous methods are advantageous because they maintain the natural state of tissue and preserve antigens and fluorescent reporters that can be lost using hydrophobic solvents and alcohols. Indeed, aqueous and hyperhydration methods such as Clarity, Sca*l*e^3,4^, ClearT^2^^5^, SeeDB^6^, FRUIT^7^, CUBIC^8–10^, and EZ Clear^12^ are among those most widely used because they are simple, non-toxic, compatible with *in vivo* tracers, and can preserve fluorescent reporters^1^. Conversely, solvent based methods such as iDISCO+^11^ and have been broadly implemented because they are robust and generally more effective on widely varying tissue types. However, as noted, solvent based techniques by and large use solvents that can be toxic and difficult to work with, and can diminish fluorescent reporter protein signal and protein antigenicity.

Tissue types vary on how amenable they are to optical clearing and 3D imaging. For instance, tough fibrous tissue, such as muscles, are very difficult to optically clear. Moreover, the autofluorescence naturally exhibited by tissue remains a major challenge in 3D imaging, as it limits the ability to detect signals from fluorescently labeled structures^1^. This presents difficulties for multiplex 3D imaging of organs such as the kidney that exhibits notoriously high background noise, or the heart that is both fibrous and has significant autofluorescence. Indeed, interrogating human biopsies of such organs are markedly challenging as they are tougher and exhibit even higher autofluorescence than analogous tissues of commonly used small mammalian animal models.

Here, we developed Atacama Clear (ATC, named for the clear skies of the Atacama Desert), an aqueous optical clearing protocol designed specifically to address the difficulties associated with imaging tough fibrous tissues and those that naturally exhibit high background noise. We demonstrate ATC potentiates the clearing capacity of all tested optical clearing chemicals and generates signal-to-noise ratios (SNRs) that are up to 200% that of SNR obtained with widely used 3D imaging methodologies. Strikingly, ATC also generated increased fluorescent reporter protein resolution, which is instead diminished with commonly used 3D imaging techniques. The advantageous properties of ATC facilitated the imaging of poorly recognized antigens, up to 10-fold dilution of antibody concentration, and the multiplex interrogation of highly autofluorescent and fibrous tissues, including the heart, kidney, and human biopsies, with cellular resolution. Taken together, these studies validate the use of ATC for complex 3D imaging of challenging basic and human samples with superior resolution.

## RESULTS

### Optimization of optical clearing protocols

Although a plethora of optical clearing protocols have been developed over the years^1,2^, to the best of our knowledge, an aqueous technique has yet to be optimized for imaging studies that are challenged due to inherently high degrees of background noise. To fill this technical gap, we sought to maximize SNRs in optical clearing studies, particularly for organs such as the kidney that exhibits high background autofluorescence (AF), and those such as the heart that demonstrate high AF and are also difficult to clear due to a robust cytoskeletal network. Indeed, as shown in Fig. 1A with the heart, kidney and spleen imaged in the same field of view, the kidney and heart exhibit relatively high levels of AF.

**Figure 1.**
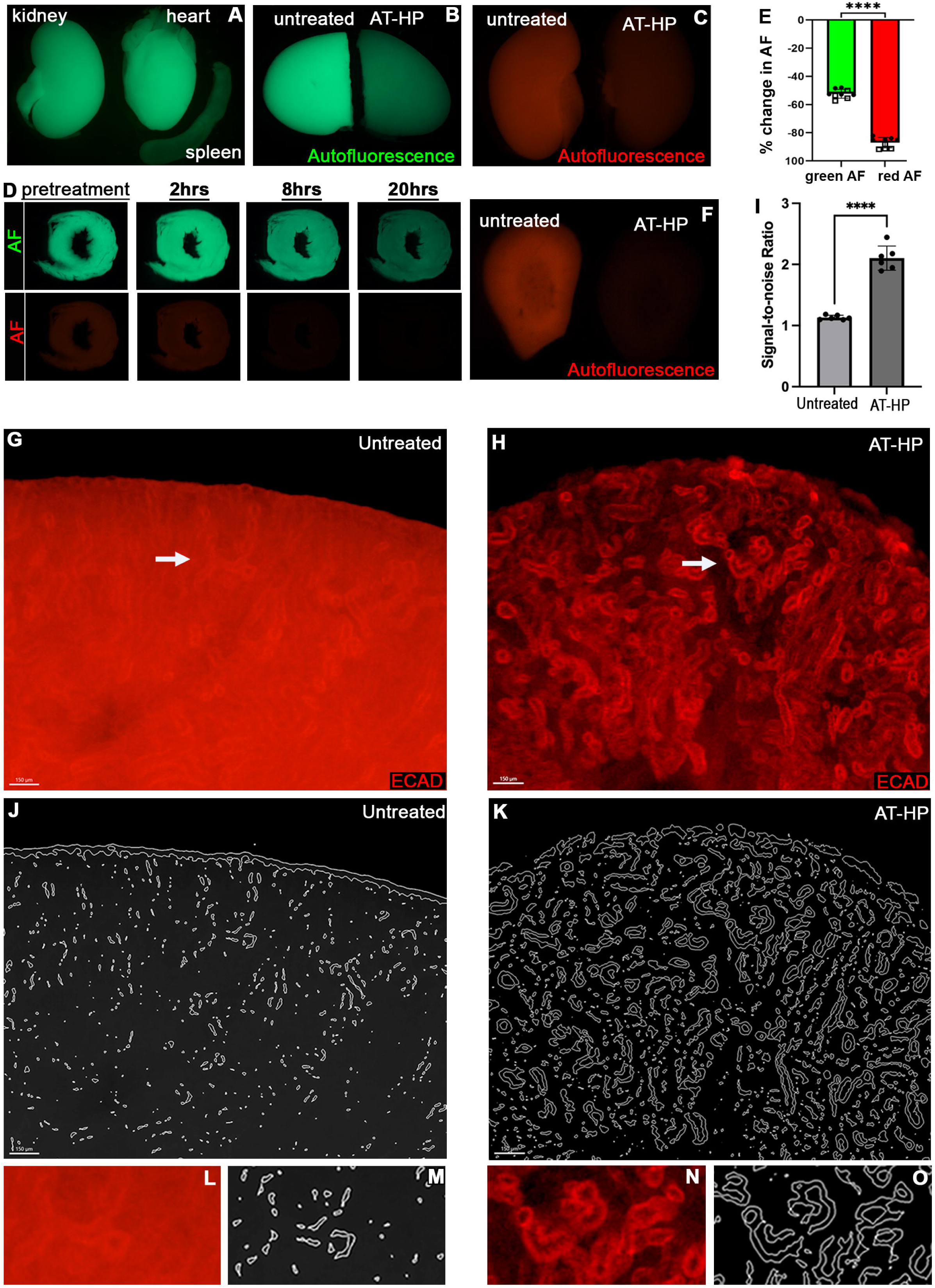
AT-HP provides superior imaging resolution and facilitates use of auto-segmentation workflows with low-tech microscopy. **A)** Stereo fluorescence microscopy of the green autofluorescence (AF) naturally exhibited by the murine kidney, heart and spleen, imaged side-by-side. **B)** Green AF exhibited by a single kidney cut in two, with one half left untreated and the adjacent half treated with AT-HP. **C)** Red AF exhibited by control untreated and AT-HP treated kidneys. **D)** Time dependent quenching of green and red AF in 1 mm thick murine heart sections. **E)** Quantification of the decrease in green and red AF in the kidney and the heart with AT-HP treatment (p<.0001). **F-H)** Stereo fluorescence microscopy imaging of adjacent kidney sections treated with and without AT-HP (F). Subsequent IF and stereo fluorescence microscopy imaging of ECAD to mark the nephrogenic renal tubules (G and H). Arrows in G and H are analyzed at high magnification in L and N, respectively **I)** Quantification of signal-to-noise ratios (SNRs) in untreated and AT-HP treated IF. **J, K)** Artificial intelligence (AI) based auto-segmentation of ECAD staining in untreated (J) and AT-HP treated tissues. **L-O)** High magnification analyses of IF staining and auto segmentation of renal tubules in untreated (L and M) and AT-HP treated (N and O) treated tissues.

Blood is a known source of AF and deterrent of optical clearing. This can be readily seen, for instance, in perfused murine hearts where regions of the organ flush with blood exhibit the highest AF (Supplemental Fig. 1A, white arrow, atria, region of low AF; black arrowhead, ventricles, region of high AF; inset, bright field image), and are also the most resistant to optical clearing (Supplemental Fig. 1B, cleared heart tissue; 1C, transparency of cleared heart region 1 vs. 2). Thus, a blood removal method would be beneficial for mitigating background noise and augmenting optical clearing. As shown in these examples, however, perfusion is often variable and incomplete. Notably, Hydrogen peroxide (H_2_O_2_) has been widely used to remove blood, as well as for quenching autofluorescence. However, it is well established that H_2_O_2_ damages tissue as catalase converts H_2_O_2_ to water and oxygen, and the accumulation of oxygenation causes tissue tearing and AF of damaged tissue regions^13^. Indeed, tissue tearing was readily observed when heart or kidney sections were treated with standard H_2_O_2_ treatment (Supplemental Fig. 1D and 1E, arrow heads mark sites of tissue tearing, arrows mark oxygenation within tissue). Tissue regions damaged by H_2_O_2_ oxygenation exhibited increased AF, as demonstrated in kidney sections before and after treatment with standard H_2_O_2_ (Supplemental Fig. 1F, before treatment; 1G, after treatment, arrows point to regions of tissue damage that increased in AF).

To address this challenge, we aimed to develop an alternative blood removal and AF quenching technology. Since oxygenation corresponded with tissue damage and increased AF, we screened eight putative reversible inhibitors of catalase to moderate oxygen production while maintaining blood removal (Supplemental Fig. 1H). Sodium azide (NaN_3_) used with H_2_O_2_, termed Atacama-Hydrogen Peroxide (AT-HP), precluded visible oxygenation (Supplemental Fig. 1H, effervescing observed in all other conditions, but not with NaN_3_). In contrast to standard protocols, treatment of 1 mm heart sections and whole kidneys showed that AT-HP fully decolorized tissue without causing tissue degradation (Supplemental Fig. 1I-K, kidney; 1L-N, heart). Critically, AT-HP significantly reduced tissue AF, as illustrated in kidneys divided into two equal halves, with one half left untreated and the second treated with AT-HP (Fig. 1B, green AF of untreated and treated kidney halves imaged in the same field of view). AT-HP progressively diminished AF in a time dependent manner (Supplemental Fig. 1O-Q), resulting in a compelling difference in background noise between treated and untreated kidney tissue, in both green (Fig. 1B) and red AF (Fig. 1C). Time dependent quenching of green and red AF was validated in 1 mm thick heart sections (Fig. 1D and Supplemental Fig. 1R-T). Treatment for up to 72 hr with AT-HP facilitated an average of 52% decrease in green AF and >85% decrease in red AF of kidney and heart tissue (Fig. 1E, mean reduction in green AF = 52% ± 2.9%; mean reduction in red AF = 87% ± 3.6%; p<0.0001). Direct comparisons of adjacent 500 µm sections cut from an individual heart or kidney that were treated for 24 hr with AT-HP or CUBIC-1, a well published decolorizing method (Supplemental Fig. 1U-X), showed AT-HP more efficaciously decolorized tissue (Supplemental Fig 1U, W) and better-preserved tissue morphology (Supplemental Fig. 1V, X). Although sections were not torn by CUBIC-1, they were enlarged and distorted in shape. By contrast, ATC sections maintained their normal morphology.

We hypothesized that the quenching of AF and decreased background noise, along with the high tissue preservation obtained with AT-HP would facilitate superior immunofluorescence (IF) signals with significantly higher SNRs. To test this hypothesis, we assayed the kidney which is notoriously difficult to image due to its high AF. Specifically, we chose to image the renal nephrons, the most autofluorescent segment of the kidney and among the highest of any tissue. To perform direct comparisons, individual unperfused adult murine kidneys were cut at 0.5 mm thickness, and resultant adjacent sections were treated with or without AT-HP (Fig. 1F) and then IF performed in the same receptacle for ECAD to mark the nephrogenic renal tubules (Fig. 1G-O). Analyses by simple, stereo fluorescence microscopy showed a striking difference between IF staining in untreated (Fig. 1G) and AT-HP treated (Fig. 1H) tissues. Despite the low-tech (e.g. lack of optical sectioning), diffuse illumination and low magnification of stereo microscopy, renal tubules were robustly detected in AT-HP kidney sections. By contrast, renal tubules were not readily detectable in standard IF of untreated adjacent sections. Indeed, AT-HP treated tissues exhibited SNRs that were 97% greater than those of untreated tissues (Fig. 1I). These data were further corroborated using artificial intelligence (AI) based auto-segmentation, which readily detected and segmented the renal tubules in AT-HP treated tissues (Fig. 1K), but not in untreated standard IF tissues (Fig. 1J). High magnification analyses validated the increase in resolution, with AT-HP enabling visualization of separate, distinct points in adjacent renal tubules, as well as the luminal area of individual tubules (Fig.1N, O, high magnification of region marked by arrow in 1H). This level of cellular resolution was not achieved with standard IF (Fig. 1L, M, high magnification of region marked by arrow in 1G).

To further substantiate the advanced resolution provided by Atacama based imaging, we next compared IF with AT-HP versus IF with well published and highly used 3D imaging methods. For these studies, we performed IF without refractive index matching, or clearing, so as to provide direct comparisons of IF signal detection, including imaging tissue in the same field of view. Notably, hydrogen peroxide has been included as a bleaching step in several leading methods^11,14,15^, including DISCO-based techniques. iDISCO+ for instance not only includes a bleaching step, but also glycine plus heparin treatment to reduce background in IF. Thus, we began our assessment by comparing IF with AT-HP versus IF with iDISCO+. We chose to interrogate the heart, which as aforementioned, is difficult to image due to its tough fibrous makeup and high endogenous background AF. Specifically, we performed imaging of the sinoatrial node (SAN), the pacemaker of the heart^16^ (Fig. 2A-E). It is well established that cells comprising the SAN express the HCN4 pacemaker ion channel^17,18^. However, whole tissue imaging of the SAN can show diffuse levels of HCN expression. Indeed, low magnification stereo fluorescence microscopy of whole atria immunolabeled for HCN4 by iDISCO+ exhibited faintly localized HCN4 signal at the terminal crest, the known anatomical site of the cardiac pacemaker (Fig. 2A, low magnification; 2B and 2D, high magnification of SAN). By contrast, whole atria imaged using AT-HP exhibited striking HCN4 signal that distinctly marked the SAN along the terminal crest (Fig. 2A, AT-HP treated atria imaged side-by-side to iDISCO+ treated atria; Fig. 2C and 2E, high magnification of AT-HP tissue).

**Figure 2.**
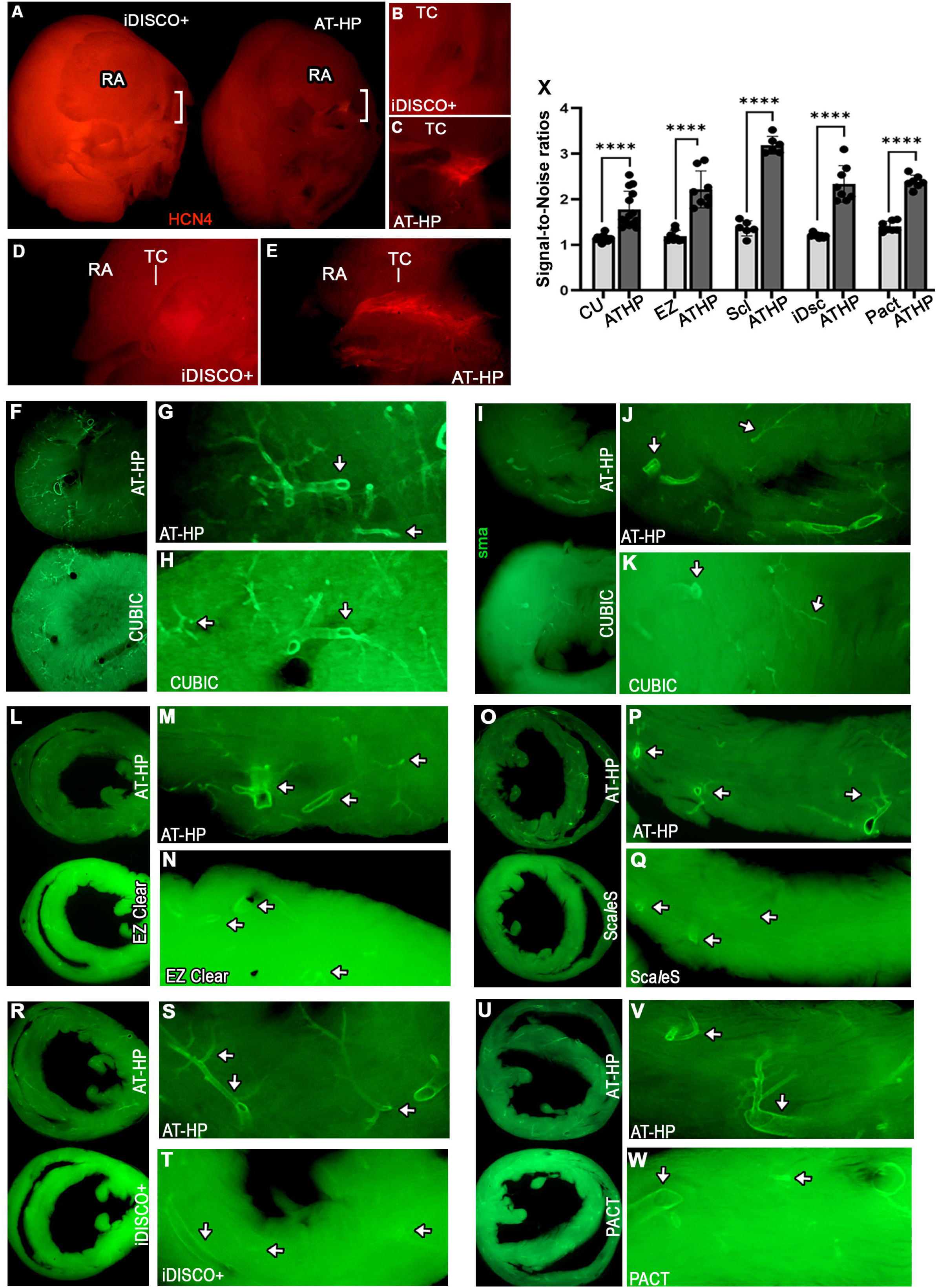
AT-HP provides superior IF resolution with significantly higher SNRs as compared to widely used 3D imaging technologies. **A-E**) Low magnification stereo fluorescence microscopy of IF performed on whole atria for the HCN4 pacemaker ion channel (HCN4, red) that marks the sinoatrial node (SAN). Atria interrogated using iDISCO+ or AT-HP were imaged side-by-side for comparison (A). High magnification analyses of bracketed regions in A showed a distinct HCN4+ node at the terminal crest (TC) of hearts imaged with AT-HP (C, coronal view; E, lateral view), which was not readily discernible in hearts imaged by iDISCO+ (B, coronal view; D, lateral view). **F-X)** Stereo fluorescence microscopy of adjacent sections cut from a single kidney or heart that were stained for arterial blood vessels (SMA, green) using AT-HP or comparative 3D imaging method, and then imaged side-by-side in the same field of view. Comparison shown to CUBIC-1 (F-H, kidney; I-k, heart), EZ Clear (L-N, heart), Sca*l*eS (O-Q, heart), iDISCO+ (R-T, heart), and PACT (U-W, heart). (X) Quantification of IF SNRs.

Notably, AT-HP exhibited drastically lower background AF in the myocardium of the ventricle (V) and atria (RA) as compared to iDISCO+ treated tissue (Fig. 2A) that were imaged side-by-side. This robust difference in background AF was corroborated in adjacent ventricular sections cut from a single heart, which showed AT-HP tissues exhibit significantly lower AF than counterpart tissues treated with iDISCO+ (Supplementary Fig. 2A-C).

AT-HP was also discovered to outperform other tested leading aqueous and solvent based techniques including CUBIC, EZ Clear, Sca*l*eS, and hydrogel-based passive Clarity (PACT). Again, to perform direct comparisons, IF was performed for the vascular bed on adjacent 0.5 mm thick sections cut from individual kidneys and hearts. IF was performed on adjacent sections with AT-HP or an alternate method, and then imaged side-by-side in the same field of view. Consistent with our earlier comparisons, AT-HP treated tissues exhibited significantly lower green and red tissue AF as compared to sister sections treated with CUBIC (Supplementary Fig. 2D, E, kidney; 2F, G, heart), EZ Clear (Supplementary Fig.2H-J, heart), Sca*l*eS (Supplementary Fig. 2K-M), and PACT (Supplementary Fig. 2N-P).

Critically, AT-HP also generated superior immunofluorescent signals compared to those obtained by these alternate techniques. Imaging of adjacent sections side-by-side in the same field of view revealed striking staining with significantly lower background in AT-HP treated tissues as compared to CUBIC (Fig. 2F-H, kidney; 2I-K, heart), EZ Clear (Fig. 2L-N, heart), Sca*l*eS (Fig. 2O-Q, heart), iDISCO+ (Fig. 2R-T, heart), and PACT (Fig. 2U-W; heart). This is further shown in enlarged micrographs of all adjacent stained sections side-by-side for all techniques (Supplemental Fig. 3). High magnification analyses (Fig. 2F-W, arrows) revealed labeled vascular structures with significantly greater resolution using AT-HP versus comparative methods, with AT-HP providing SNRs that are 57% greater than CUBIC (Figure 2X, SNR: CUBIC = 1.144 ± 0.06; AT-HP = 1.804 ± 0.39, p<.0001), 86% greater than EZ Clear (Fig. 2X, SNR: EZ Clear = 1.190 ± 0.12; AT-HP = 2.216 ± 0.40, p<.0001), 133% greater than Sca*l*eS (Fig. 2X, SNR: ScaleS= 1.36 ± 0.16; AT-HP = 3.186 ± 0.19, p<.0001), 94% greater than iDISCO+ (Fig. 2X, SNR: iDISCO+ = 1.198 ± 0.04; AT-HP = 2.33 ± 0.40, p<.0001), and 70% greater than PACT (Fig. 2X, SNR: PACT = 1.402 ± 0.11; AT-HP = 2.382 ± 0.144, p<.0001).

To date, 3D imaging protocols have, at best, retained varying levels of endogenous fluorescent reporter protein signals. Based on our results showing increased immunofluorescence signal detection and SNRs, we hypothesized that AT-HP could alleviate this technical challenge. To this end, AT-HP was used to treat heart tissues exhibiting RFP in the coronary vasculature, as well as kidney and testis tissues exhibiting Venus reporter protein in interstitial and spermatogonial stem cells, respectively^19^. AT-HP strikingly enhanced the capability to detect reporter proteins in all cases (Fig. 3). In the heart, although both RFP+ large vessels and microvasculature could be detected prior to treatment, the demarcation of these structures was difficult to differentiate from background, especially for the smaller vasculature (Fig. 3A, low magnification image; 3C and 3E, high magnification images of large vessels and the microvasculature, respectively). Imaging of the exact same tissues after AT-HP revealed markedly higher signal resolution with a 64% increase in SNRs, which was evident by the now well-defined boundaries of both large vessels and microvasculature and an abrogated background noise (Fig. 3B, 3D, 3F and 3G, RFP).

**Figure 3.**
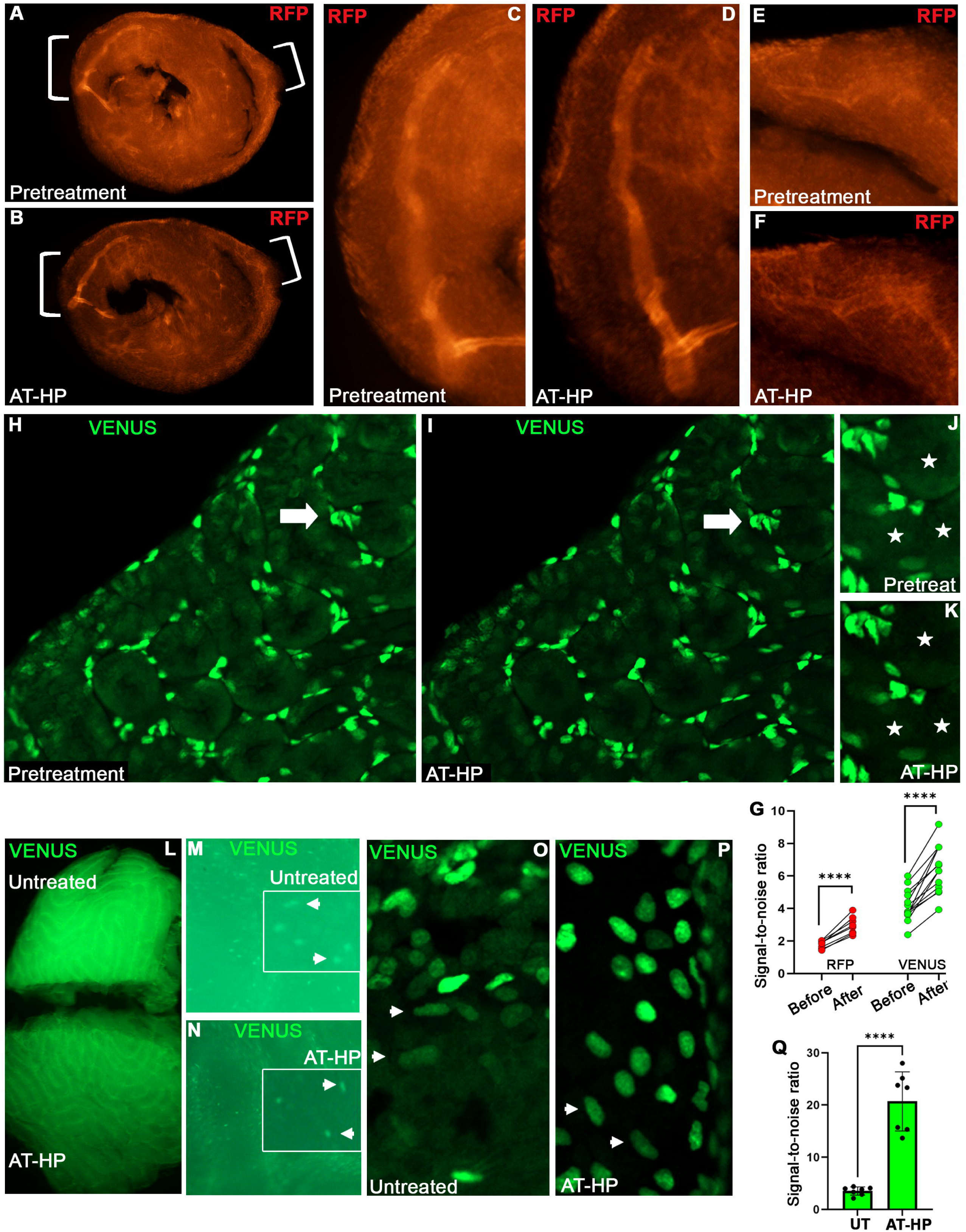
AT-HP facilitates superior imaging of fluorescent reporter proteins with increased SNRs and resolution. Imaging of transgenic murine organs expressing different fluorescent proteins with or without AT-HP treatment. **A-F)** Imaging of heart sections expressing red fluorescent protein (TdTomato) in the vasculature prior to (A, C, and E, pretreatment) or after AT-HP treatment (B, D, and F, AT-HP). Brackets in A and B indicate regions shown at high magnification in C-F. **G)** SNR levels of endogenous fluorescent reporters before and after treatment with AT-HP (p<.0001). **H-K)** Imaging of kidney sections expressing Venus green fluorescent protein in interstitial cells surrounding the renal tubules before (H and J, pretreatment) or after AT-HP treatment (I and K, AT-HP). Arrows in H and I point to the same tissue regions shown at high magnification in J and K. Asterisks mark centers of renal tubules. **L-P)** A testis expressing Venus in tubular interstitial cells was split in two, with one half left untreated and the second half treated with AT-HP (L, untreated and treated tissue shown side-by-side). Low magnification stereo fluorescence microscopy showed a marked difference in detectable Venus+ cells between untreated and AT-HP treated samples (M and N; inset shows higher magnification of Venus+ cells, arrowheads mark representative cells). O, P) High magnification imaging of Venus+ cells in untreated (O) and treated tissues (P). Arrowheads in O and P mark representative cells exhibiting low Venus+ expression in each condition. Q) SNR levels of Venus+ signal in cells of untreated (UT) or AT-HP treated testis tissue.

Analyses of kidney tissues expressing Venus corroborated our findings in the heart. Here, the Venus signal (green) exhibited by the circular interstitial cells could be directly compared to the distinct AF exhibited by the renal tubules that these cells surround. Imaging of the exact same tissue before (Fig. 3H and 3J) and after treatment (Fig. 3I and 3K) showed AT-HP nearly abolished renal tubular AF, while Venus reporter signal remained unchanged. This was reflected in an 55% increase in SNRs after AT-HP treatment (Fig. 3G, Venus).

AT-HP was next used to facilitate detection of Venus+ interstitial and spermatogonial stem cells in testis, an organ that also exhibits marked tissue AF^20^. One half of a bisected testis was treated with AT-HP and the other left untreated (Fig. 3L). Low magnification stereo fluorescence microscopy showed Venus+ cells were more visible in AT-HP treated tissues compared to untreated tissues (Fig. 3M, untreated tissue; 3N, AT-HP treated tissue; insets highlight Venus+ signals). These findings were corroborated by high magnification imaging of Venus+ cells (Fig. 3O, untreated tissue; 3P, AT-HP treated tissue). In untreated tissues, bright Venus+ cells were distinguishable, whereas cells with low Venus+ signal were difficult to distinguish from the background tissue AF (see arrows in Fig. 3O). Conversely, background tissue AF was abolished in AT-HP treated tissue, consistent with our findings observed in the kidney. With autofluorescence rendered negligible, AT-HP treatment readily facilitated not only the detection of bright Venus+ cells, but also the distinct visualization of cells exhibiting lower Venus+ signal (see arrows in Fig. 3P). AT-HP treated testis tissues exhibited Venus+ SNRs that were a striking 5.8x higher than untreated tissues (Fig. 3Q).

In addition to these advantageous properties, we also discovered that the superior SNRs facilitated by AT-HP enabled the use of up to 10-fold lower antibody concentrations for immunofluorescence. Specifically, we performed immunofluorescence of heart, kidney, and testis tissue that exhibit high background with widely used and well-documented antibodies (Fig. 4). An individual 0.5 mm thick tissue section of each organ was halved, with one half left untreated and the second half treated with AT-HP, and then immunofluorescence performed on both halves in the same receptacle. Coronary arteries (Fig. 4A, B, heart; SMA, red), nephrogenic renal tubules (Fig. 4E, F, kidney; ECAD, red), and testicular tubules (Fig. 4I, J, testis; MCAM, red) could be detected in both untreated and AT-HP treated tissues at standard concentrations. Next, we performed immunofluorescence using 2-, 4-, and 10-fold lower primary antibody concentrations (Fig. 4C, D, G, H, K, and L, and Supplementary Fig. 4). AT-HP treated tissues continued to exhibit high SNRs at all concentrations, whereas IF signal was diminished in untreated tissues at 4x lower antibody concentrations and below. Indeed, AT-HP treated tissues still exhibited robust immunofluorescence signals at 10-fold lower concentrations (Fig. 4D, heart; H, kidney; L, testis), whereas untreated tissues lacked detectable signals at this concentration (Fig. 4C, heart; 4G, kidney; 4K, testis).

**Figure 4.**
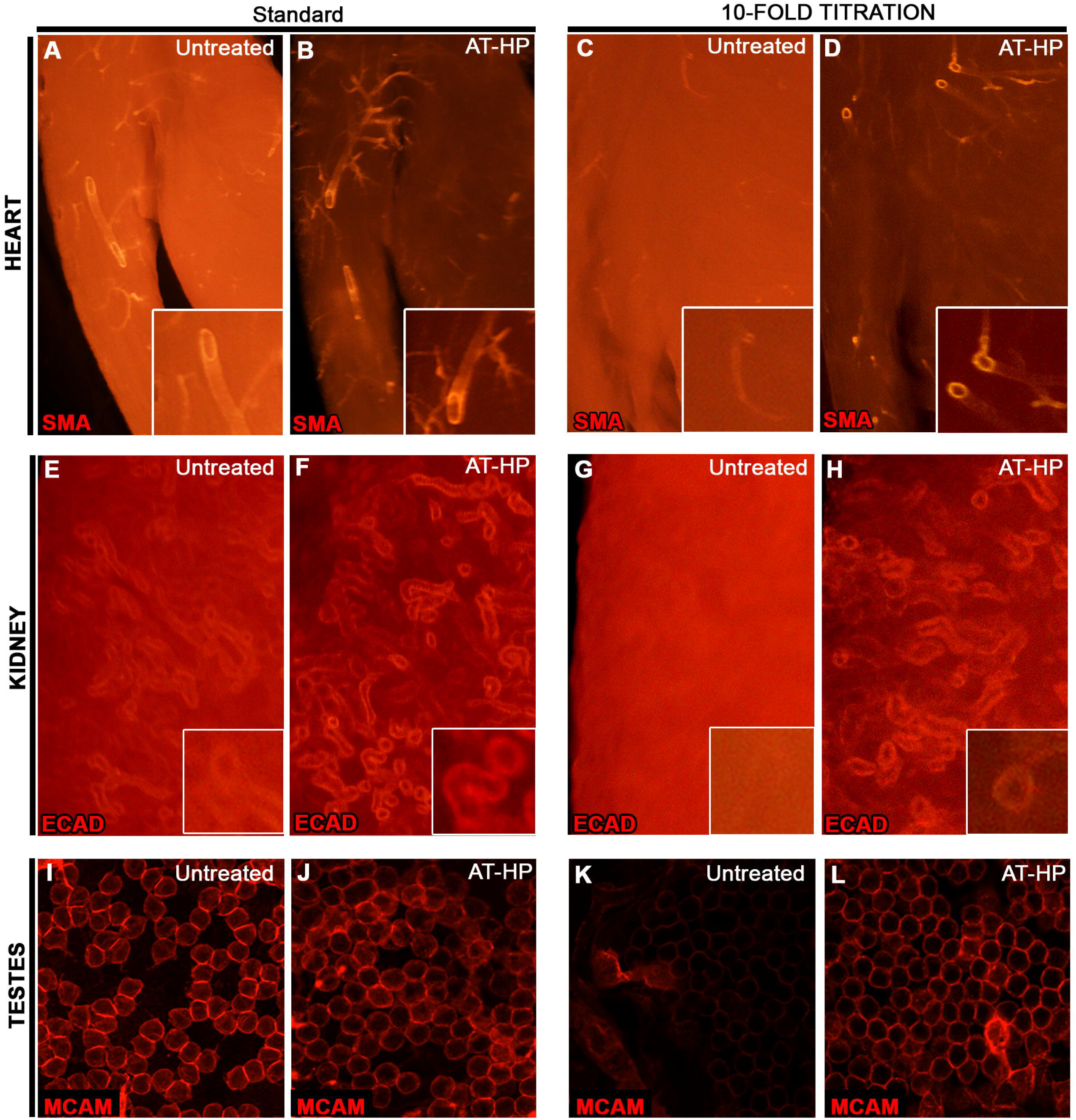
AT-HP facilitates the use of up to 10-fold lower antibody concentrations in IF. IF was performed using highly published antibodies on heart, kidney and testis tissues that exhibit high AF. **A-D)** IF for the arterial vasculature (SMA, red) using standard published antibody concentrations (A and B) or 10-fold lower concentration (C and D) on an individual halved heart section, where one half was left untreated and the other treated with AT-HP. **E-H)** IF for the renal collecting tubules (ECAD, red) on a halved kidney section, where a half was left untreated and the other treated with AT-HP, using standard published antibody concentrations (E and F) or 10-fold lower concentration (G and H). **I-L)** IF for germ cells (MCAM, red) on testis sections cut from the same testis that were treated without or with AT-HP using standard published antibody concentrations (I and J) or 10-fold lower concentration (K and L).

Notably, consistent with our initial hypothesis, we also discovered that AT-HP significantly augments optical clearing. To test this, adjacent 0.5mm sections from non-perfused kidneys were left untreated or treated with AT-HP, and then both sections were immersed in refractive index matching optical clearing solutions from published 3D imaging techniques, and imaged side-by-side (Fig. 5A-I). Incubation with EZ view (EZ Clear protocol) for instance showed a striking difference with and without AT-HP treatment. Live imaging revealed almost instantaneous clearing of AT-HP tissues, whereas untreated tissues took longer to begin clearing and never reached the same level of transparency (Fig. 5A, EZ View and Supplementary Movie 1). Similar findings were shown by live imaging of AT-HP treated tissues with CUBIC-2 (CUBIC protocol, Fig. 5B and Supplementary Movie 2), MACS-R2 (MACS protocol, Fig. 5C and Supplementary Movie 3) and FOCM (FOCM protocol, Figure 5D and Supplementary Movie 4), with AT-HP tissues exhibiting faster and more thorough optical clearing. Indeed, treatment with AT-HP increased the optical clearing capacity of all tested refractive index matching solutions (Fig. 5K), giving a striking increase in transparency of over 600% for CUBIC-2 (Fig. 5K, grid visibility: AT-HP = .822 ± 0.213; Untreated = 0.112 ± 0.074), 280% increase for EZ View (Fig. 5K, grid visibility: AT-HP = 0.838 ± 0.052; Untreated = 0.218 ± 0.142), 470% increase for MACS-R2 (Fig. 5K, grid visibility: AT-HP = 0.611 ± 0.122; Untreated = 0.106 ± 0.043), 200% increase for FOCM^21^ (Fig. 5K, grid visibility: AT-HP = 0.415 ± 0.138; Untreated = 0.134 ± 0.065), 320% increase for Sca*l*eA2 (Sca*l*eS protocol^4^; Fig. 5K, grid visibility: AT-HP = 0.293 ± 0.072; Untreated = 0.069 ± 0.035), 330% increase for ClearT (ClearT^2^ protocol^5,22^; Fig. 5K, grid visibility: AT-HP = 0.570 ± 0.133; Untreated = 0.130 ± 0.054), 260% increase for glycerol^23^ (Fig. 5K, grid visibility: AT-HP = 0.719 ± 0.112; Untreated = 0.197 ± 0.066), 210% increase for SeeDB (SeeDB protocol^6^; Fig. 5K. grid visibility: AT-HP = 0.408 ± 0.099; Untreated = 0.130 ± 0.056), and 90% increase for sRIMS (PACT protocol^24^; Fig. 5K, grid visibility: AT-HP 0.379 ± 0.027; Untreated = 0.197 ± 0.044). Given these findings, we termed the combined use of AT-HP with a refractive index matching solution Atacama Clear (ATC), for instance AT-HP + EZ View, termed ATC-EZ View. Strikingly, ATC-EZ View was shown to optically clear whole adult murine hearts in 3 hours (Figure 5J), without need of chemical delipidation, whereas EZ View studies report 24 hours of refractive index matching time for the heart, with partial optical clearing. This marks an 8-fold decrease in optical clearing time.

**Figure 5.**
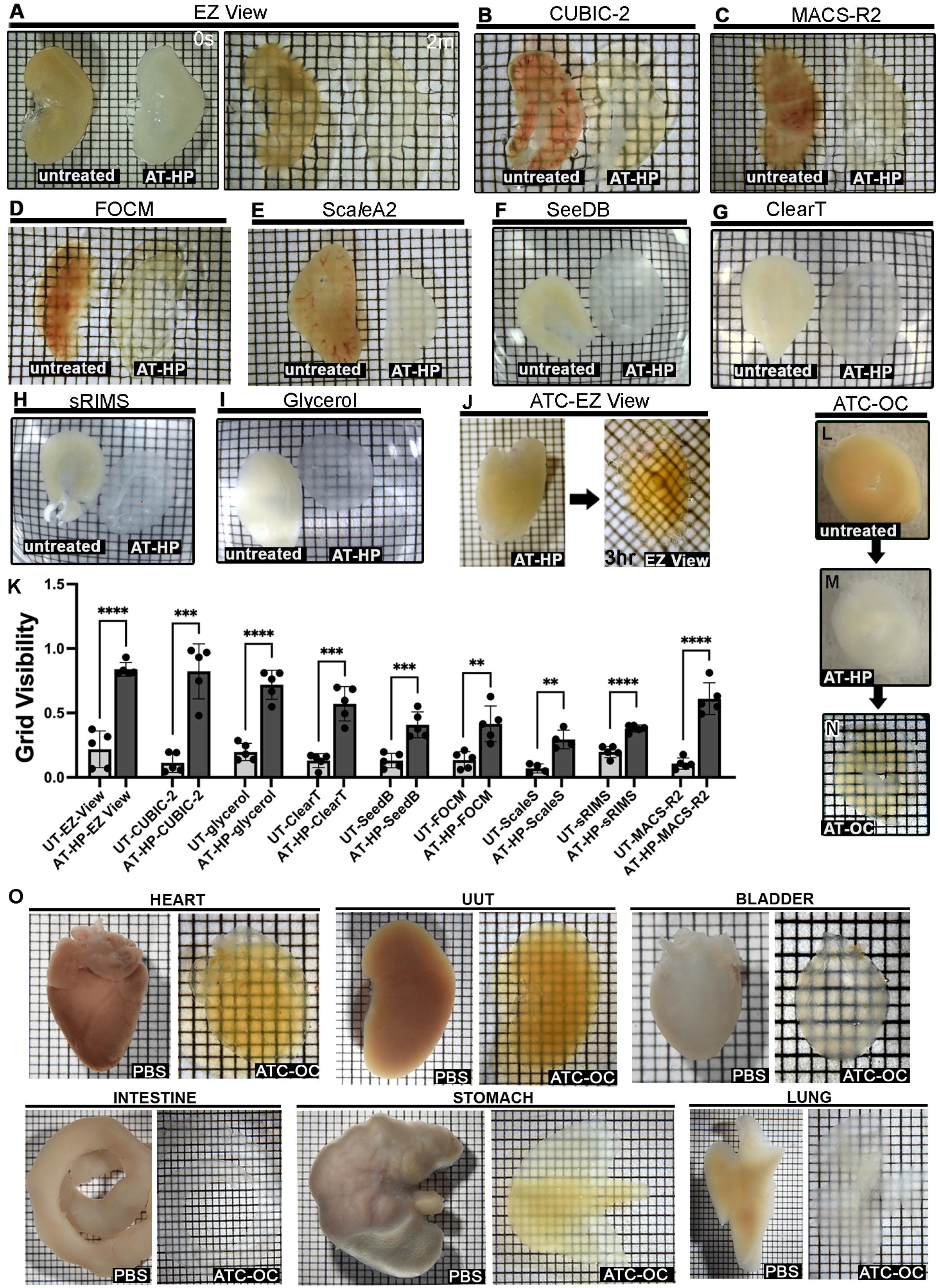
Augmented optical clearing with AT-HP. **A-I**) 0.5 mm thick sections obtained from a single kidney were left untreated or treated with AT-HP, and then imaged in the same field of view after treatment with refractive index matching, optical clearing solutions used in published and widely used 3D imaging clearing protocols, including EZ View (A, left, time 0 right before treatment; right, 2 min after treatment; also see Supplemental Movie 1), CUBIC-2 (B, also see Supplemental Movie 2), MACS-R2 (C, also see Supplemental Movie 3), FOCM (D, also see Supplemental Movie 4), Sca*l*eS (E), SeeDB (F), ClearT (G), sRIMS (H), and glycerol (I). **K)** Quantification of percentage increase in optical clearing capacity of reagents after tissue treatment with AT-HP. **J)** Optical clearing of whole murine heart after 3 hrs with ATC-EZ View. **L-N)** Representative workflow of ATC optical clearing, showing an unperfused 0.5 mm thick heart section (L) being first treated with AT-HP (M) and subsequently cleared with ATC-OC (N). **O**) Representative ATC-OC optical clearing of whole murine organs, including heart, upper urinary tract (UUT), bladder, intestine, and stomach and lung.

Based on our data noting high AF with use of numerous protocols, we continued our protocol development. For instance, we imaged the AF of adjacent kidney sections before and after treatment with formamide, quadrol, or m-Xylylenediamin (MXDA), active reagents utilized in ClearT^2^, CUBIC, and MACS, respectively (Supplemental Fig. 5A-C). Treatment with formamide resulted in a decrease in AF (Supplemental Fig. 5A, representative images before and after treatment; Supplemental Fig. 5D). By contrast, treatment with 20%, 40% or 50% MXDA increased kidney AF (Supplemental Fig. 5B, images before and after treatment; and Supplemental Fig. 5D). Moreover, 20% quadrol, 40% quadrol, and CUBIC 1 reagent (25% quadrol, 25% urea, and 15% triton-x) increased kidney AF (Supplemental Fig. 5C, images before and after treatment; and Supplemental Fig. 5D). Putative increases in non-specific signal were reported in the development of CUBIC^8^. Based on these results, protocol development was continued using formamide. It was hypothesized that formamide combined with other refractive-indexed matched reagents would provide greater optical clearing capabilities. Glycerol was chosen for the following beneficial properties: a) it can clear tissue with a refractive index that closely matches biological tissues; b) it is hydrophilic, precluding the need for dehydration with hydrophobic solvents and alcohols that compromise membrane and fluorescent reporter protein integrity; c) it has excellent optical properties, including the preservation of fluorescence, and does not increase AF; and d) it is non-toxic, and simple to integrate. To test our hypothesis, we used murine heart sections due to their resistance to optical clearing. Non-perfused hearts were sectioned at varying thicknesses and then treated either with 25% formamide + 75% glycerol solution, which we termed Atacama Clear optical clearing (AT-OC) solution, or ClearT^2^ that is comprised of 50% formamide and 20% PEG (Supplemental Fig. 5E). ClearT^2^ failed to clear heart sections thicker than 500 µm. By contrast, AT-OC readily cleared heart sections of 850 µm thickness and exhibited more than a 2-fold greater clearing capacity than ClearT^2^ (Supplemental Fig. 5F). Moreover, 1 mm thick sections of perfused hearts that lacked blood were readily made transparent with AT-OC (Supplemental Fig. 5G).

Thus, we hypothesized that sequential use of AT-HP followed by AT-OC, termed ATC-OC (Fig. 5L-N), would facilitate efficacious tissue optical clearing and 3D imaging of challenging tissue types. To test this, the capacity of ATC-OC to optically clear whole murine organs was assessed. ATC-OC cleared intact muscular organs including the heart, upper urinary tract (UUT), bladder, intestine, and stomach, as well soft tissue organs, such as the lungs (Fig. 5O). While refractive index matching required several days, AF remained quenched, and moreover, measurement of kidney or heart tissue before and after treatment showed ATC-OC does not cause tissue deformation, with high preservation of morphology and size (Supplemental Fig. 5H and I).

Taken together, our protocol optimization studies established Atacama technology provides superior immunofluorescence signals with significantly higher SNRs than standard IF and leading technologies; facilitates the use of auto-segmentation workflows with low tech microscopy; enables IF with up to 10-fold lower antibody concentration; significantly increases the speed and optical clearing capacity of refractive index matching solutions to render tissue transparent; facilitates clearing of whole muscular organs; and increases the detection of endogenous fluorescent reporters, an aspect that has to date been lacking in 3D imaging techniques.

### Multiplex 3D imaging with ATC-OC

We next tested our hypothesis that the advantageous properties of ATC-OC facilitates 3D multiplex imaging of challenging tissue types. Specifically, we assayed the kidney which is notoriously difficult to image due to its high AF, particularly in the green field, which limits the ability to simultaneously visualize different organ compartments by multiplex immunofluorescence. Consequently, we chose to perform imaging of the different tissue layers of the glomerulus, which is furthermore demanding due to the high tissue preservation required to maintain the delicate architecture of this tissue^25^. As can be seen in Fig. 6A-D and Supplemental Movie 5 (3D view), the podocytes (Podocalyxin, red), mesangium (Pdgfr-β, blue) and endothelium (Endomucin, green) of the murine glomerulus were readily distinguished using ATC-OC^26,27^.

**Figure 6.**
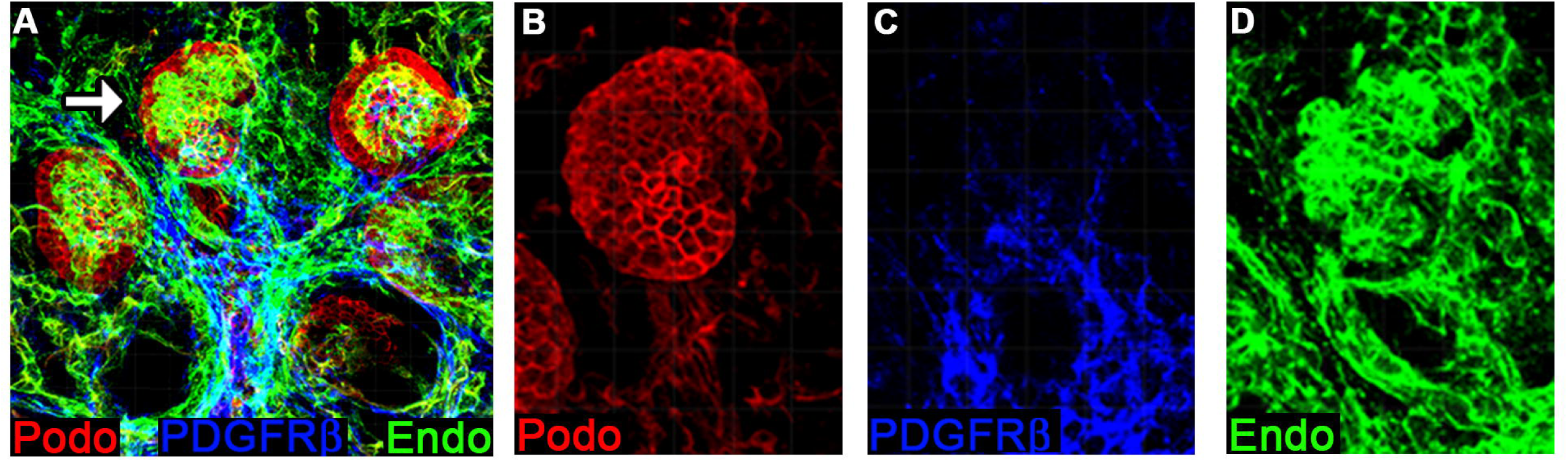
ATC facilitates multiplex IF imaging of murine renal tissue. ATC was used to perform 3D multiplex imaging of murine tissues that exhibit notoriously high AF. **A-D)** Murine renal tissues were immunolabeled for the intricate structures of the glomerulus (A, merged labeled image, arrow points to region shown at high magnification), including the podocyte (B, podocalyxin, red), mesangial (C, PDGFR-β, blue) and endothelial (D, Endomucin green) tissue compartments. All tissue regions were clearly differentiated (also see Supplemental Movie 5).

The capability of ATC-OC to detect poorly recognized and understudied global tissue structures was next assessed. Again, we chose the renal nephrons because it exhibits among the highest AF of any tissue. Indeed, the native global architecture of different nephron segments remains to be directly imaged. Thus, we used ATC-OC to image parvalbumin and ENAC that mark adjacent distal convoluted and connecting tubule nephron segments, respectively^28,29^. As shown in Fig. 7A-C and Supplemental Movie 6 (3D view), these tissue regions were distinctly visualized. Critically, we were able to interrogate the 3D structure of these tissues, such as connecting tubule length (Fig. 7B), with single cell resolution (Fig. 7C). Thus, ATC-OC facilitated 3D imaging of previously undocumented tissues, at cellular resolution.

**Figure 7.**
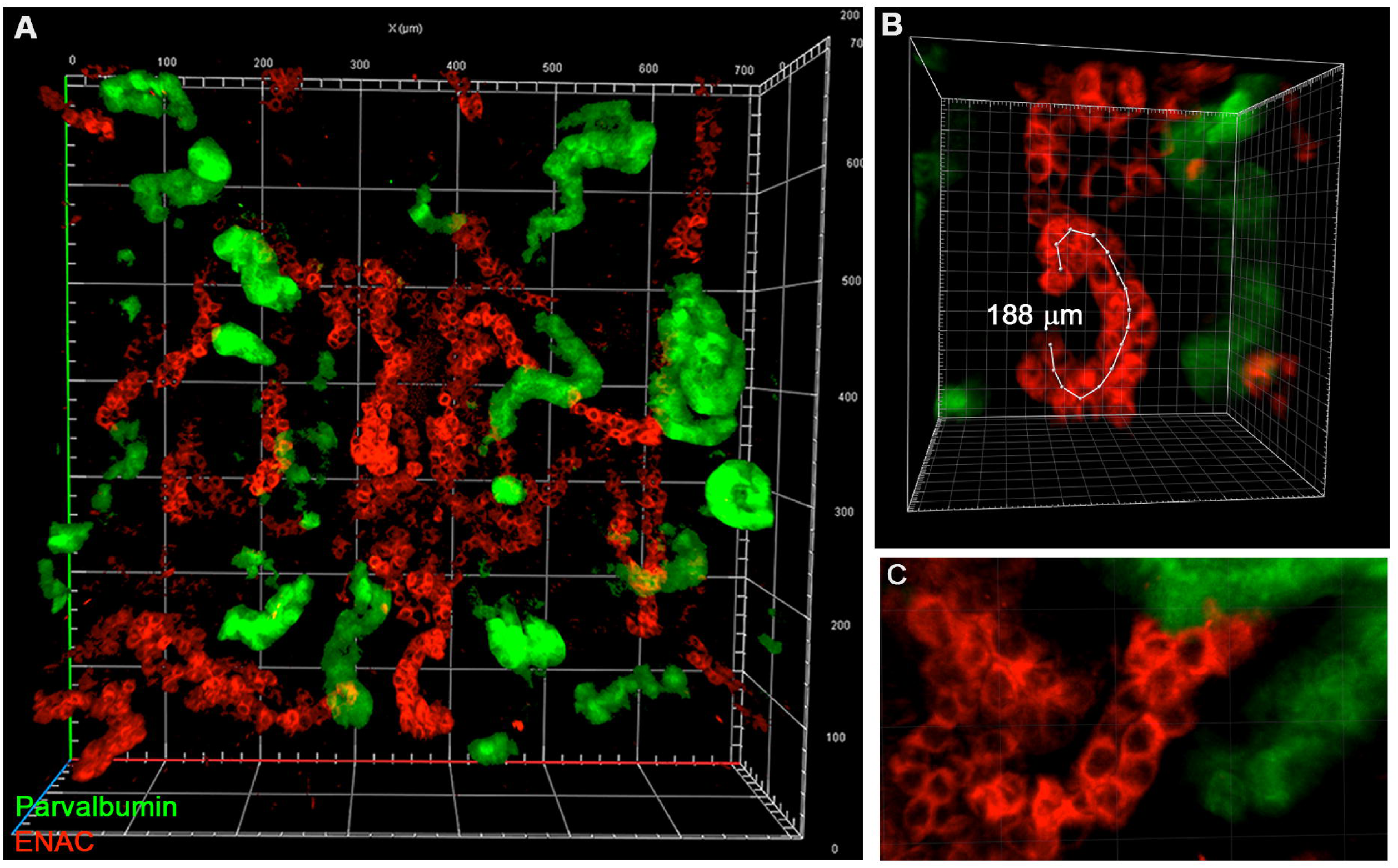
3D imaging of previously undocumented nephron tissue architecture using ATC. ATC was used to 3D image nephron segments that exhibit among the highest tissue AF. Kidney tissues were stained for ENAC (red) and Parvalbumin (green) to mark adjacent connecting and distal convoluted tubule nephron compartments, respectively. **A)** ATC readily facilitated the distinct visualization of each nephron segment, as well as their gross architectural tissue arrangement (see also Supplemental Movie 6). **B)** The high SNRs obtained using ATC facilitated direct measurements of tubule dimensions (demarked measurement of connecting tubule length in 3D = 188 µm), and **C)** provided single cell resolution.

The advantageous properties of ATC-OC to facilitate imaging of difficult to detect antigens was next corroborated in testis tissue that also exhibits high background noise (see Fig. 3). Specifically, we assayed for cKit, Stra8, and GFRα1, markers of spermatogonial stem cells and differentiating spermatogonia (Figure 8A-F)^20,30–33^. Notably, as can be seen, standard immunofluorescence signals from these markers poorly differentiate from background tissue noise (Fig. 8A, C, and E), which hinders the definitive analyses of these cell types. By contrast, imaging of these markers with ATC-OC produced robust immunofluorescence signals that distinctly differentiated from background. ATC-OC provided a striking 3x, 4.5x, and 3.5x higher SNRs for cKit, Stra8, and GFRα1, respectively (Fig. 8G). Indeed, 3D imaging of testicular tubules (Fig. 8H, MCAM, red, germ cell marker) allowed the clear visualization of spermatogonial stem cells marked by GFRα1, including A paired spermatogonia (arrows), and A aligned spermatogonia with chains of 4-cells (arrow head) and 8-cells (bracket) spermatogonial stem cells marked by GFRα1. Thus, we establish ATC-OC for the visualization of poorly recognized antigens in tissue types like the kidney and testis that are difficult to multiplex image due to their high background noise.

**Figure 8.**
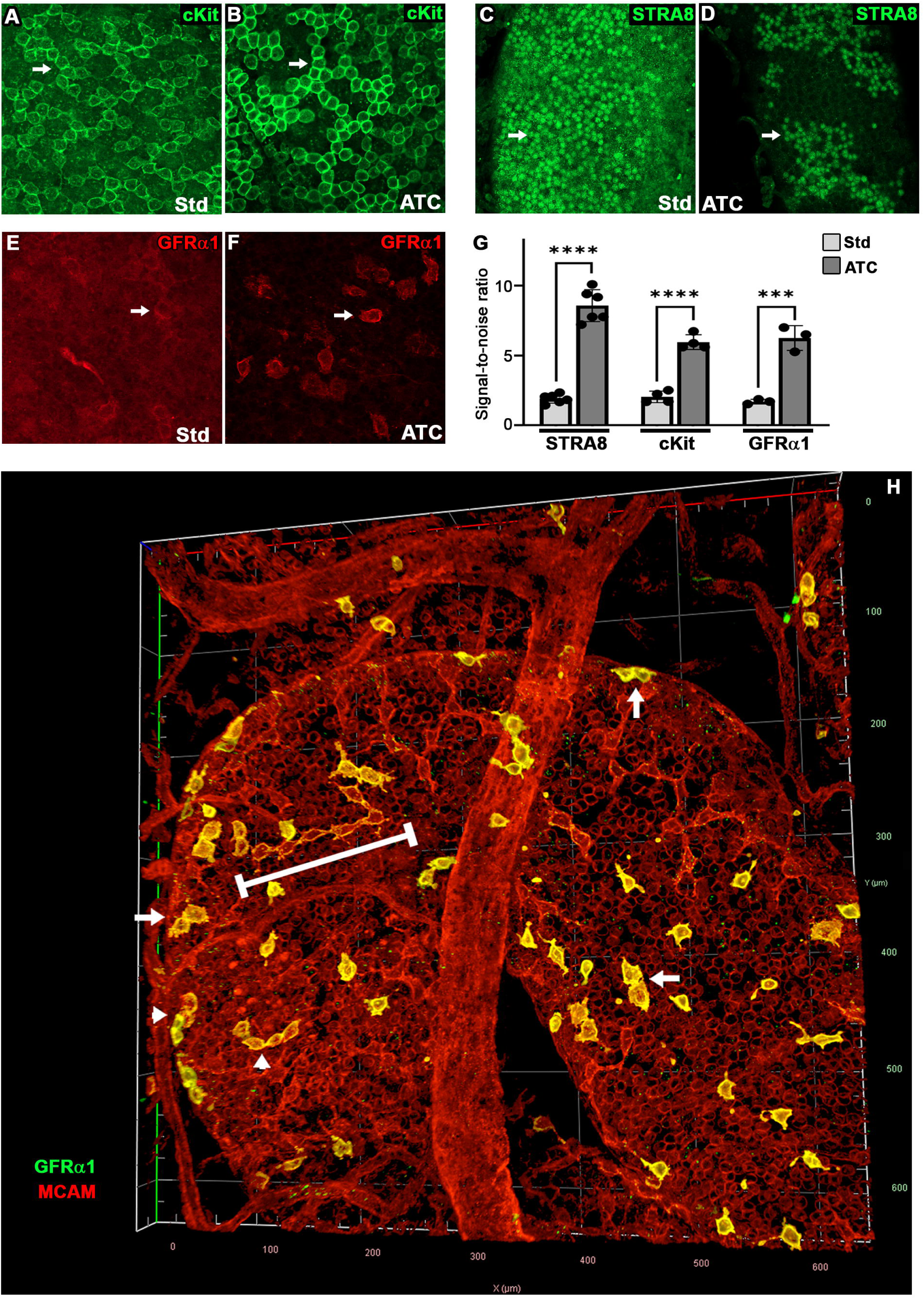
ATC facilitates superior visualization of poorly recognized antigens. ATC was used to interrogate spermatogonial stem cells markers that are poorly recognized and difficult to visualize by IF. **A-F)** Sections of the same testis were imaged by standard IF or with ATC-OC for the spermatogonia stem cell markers cKit (A and B, green), Stra8 (C and D, green), and GFR-α1 (E and F, red). **G)** SNR levels provided by standard IF or with ATC-OC. **H)** 3D imaging of testis tubules for MCAM and GFR-α1 enabled visualization of spermatogonial stem cells in A paired spermatogonia (arrows), and A aligned spermatogonia with 4 cells (arrow heads) and 8 cells (bracket) stages of differentiation.

We next examined the heart to determine the efficacy of ATC-OC to image through muscular organs, the most difficult tissues to optically clear. Murine hearts were stained for the arterial vasculature and imaged using ATC-OC (SMA, red, Fig. 9). Even using low light conditions and diffuse illumination of low magnification stereo fluorescent microscopy, the cardiac arterial bed was nonetheless readily visualized (Fig. 9A, inset). Confocal microscopy was used to determine the capability of imaging deep inside the organ. Imaging was done through the dense ventricular myocardium. Analyses of the individual slices constituting the 3D image showed no loss of labeled vessel signal intensity as deep as 1.6 mm into the heart, which was the microscopic focal working distance limit (Fig. 9B and Supplemental Fig. 6 showing all representative image slices).

**Figure 9.**
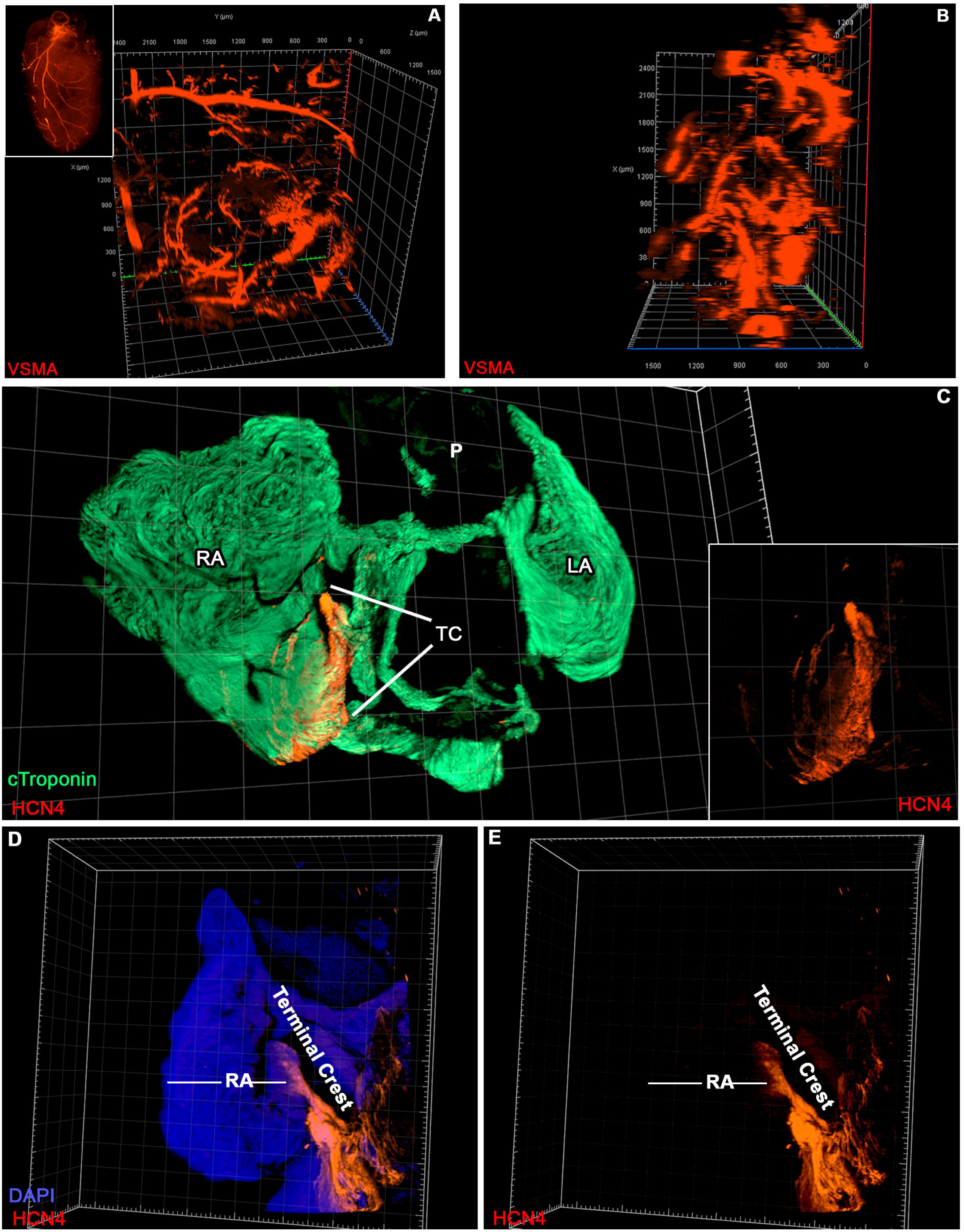
Imaging of cardiac vascular bed and SAN pacemaker of the cardiac conduction system. ATC was used to visualize the arterial bed of the heart (SMA, red). **A)** Both low magnification stereomicroscopy (inset) and confocal microscopy (A) readily facilitated imaging of cardiac arteries. **B)** With confocal microscopy ATC-OC facilitated imaging 1.6 mm deep inside the heart without loss of signal intensity (x-axis view; see also Supplementary Fig. 6, showing all the representative slices of the 3D image). **C)** 3D confocal microscopy imaging of whole atria (C) immunolabeled for both the working myocardium (cTroponin-T, green) and sinoatrial pacemaker node (HCN4 pacemaker channels, red). 3D interrogation showed that the SAN indeed localized to the terminal crest (TC) of the right atrium (C; inset shows SAN alone). **D and E)** Independent validation by staining with DAPI and HCN4 confirmed that HCN4 pacemaker channels are readily imaged and restricted to the anatomical site of the SAN, or the terminal crest of the right atria (D, merged; E, HCN4 alone). RA, right atria; LA, left atria; TC-terminal crest

To further substantiate these findings, we performed multiplex 3D imaging of the SAN (Fig. 9C-E). Consistent with our stereo microscopy analyses (Fig. 2), 3D confocal microscopy imaging with ATC-OC of whole atria immunolabeled with HCN4 and Troponin-T to mark the myocardium facilitated the localization of a robust HCN4+ SAN tissue at the terminal crest (Fig. 9C, inset HCN4+ SAN alone). Independent validation by staining solely with DAPI and HCN4 also confirmed that HCN4 is indeed highly expressed and restricted to the terminal crest of the right atrium (Fig. 9D, E). Thus, we demonstrate that ATC-OC facilitates high resolution imaging of the heart, including specialized elements of the conduction system.

We next tested the preservation of fluorescent reporter signals using ATC-OC in several experimental model systems (Fig. 10). Embryoid bodies (EBs) derived from murine stem cells with a doxycycline-inducible GFP reporter were treated with and without doxycycline, immunostained for E-cadherin (red) to label all cells, and cleared with ATC-OC. GFP was readily detected in the doxycycline-induced EBs (Fig. 10A), and is lacking in control EBs without induction (Fig. 10A, inset). GFP was also readily imaged in the ventricles of adult zebrafish hearts expressing GFP from a cardiomyocyte-specific transgenic promoter (Fig. 10B-D)^34^. Treatment with ATC-OC mitigated the AF in the zebrafish trunk, while preserving the GFP expression in the heart (Fig. 10B, before, Fig. 10C, after ATC-OC). GFP expression was detected in the ventricular myocardium, but not in the non-contracting bulbus arteriousus of the heart (Fig. 10D, inset, DAPI)^34^. We used ATC-OC to image kidneys from adult *Spry4^H2B-Venus^* transgenic mice expressing Venus fluorescent protein in renal interstitial cells^19^. As shown in Fig. 10E and F, Venus signal was efficiently preserved. Interestingly, we immunolabeled renal tissues with SMA to mark the renal arterial bed, and discovered that the renal artery contains a previously undocumented population of Venus+ interstitial cells (Fig. 10F, arrows), which are cells known to play an important role in controlling vascular tone and compliance^35^. Consistent with these data, we also established the preservation of RFP using ATC-OC (Fig. 10G and H). Murine hearts exhibiting RFP in the arterial bed were processed with ATC-OC and imaged whole using stereo fluorescence microscopy. As shown, ATC-OC preserved the robust RFP+ signal of coronary arteries (Fig. 10G), including that of smaller caliber vessels branching from the major arteries (Fig. 10H). Thus, we demonstrate the preservation of multiple fluorescent reporters, in varying model systems, using ATC-OC.

**Figure 10.**
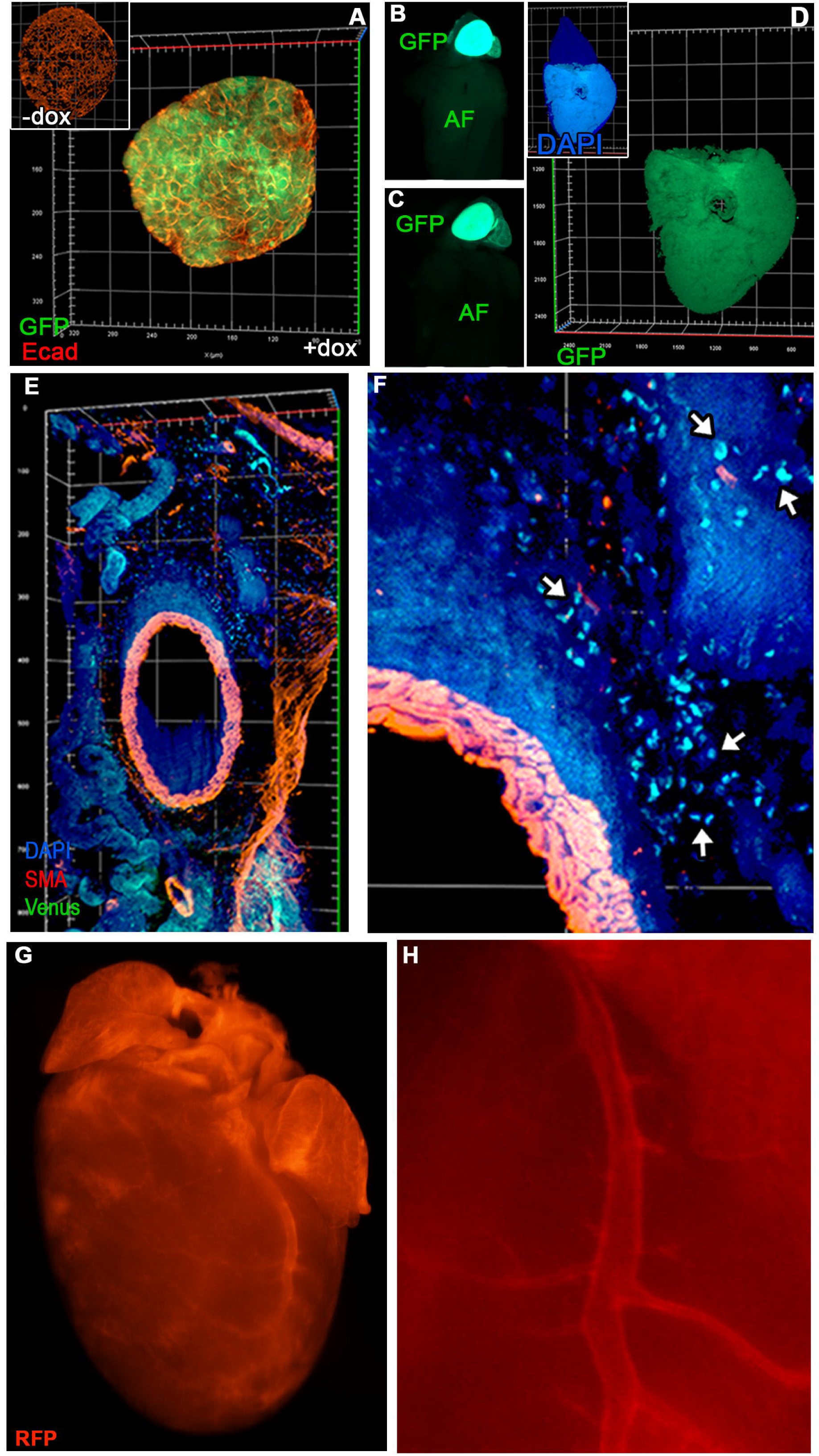
Preservation of fluorescent reporter proteins across model systems with ATC-OC. **A)** 3D imaging of intact ATC-OC cleared stem cell-derived EBs exhibiting doxycycline-inducible GFP expression and immunolabeled for E-cadherin (Inset, control organoids without doxycycline treatment). **B-D)** ATC-OC imaging of whole zebrafish exhibiting GFP expression in the ventricular myocardium of the heart showed a decrease in AF (AF) and retained GFP expression (B, before ATC-OC treatment; C, after ATC-OC treatment). 3D imaging through intact hearts validated GFP expression localized to the ventricular myocardium (D; Inset, DAPI). **E and F)** Low (E) and high (F) magnification 3D imaging of ATC cleared *Spry4^H2B-Venus^* kidneys immunolabeled for smooth muscle actin (SMA, red) revealed a Venus+ interstitial cell population localized to the renal artery (F, Venus+ interstitial cells, arrows). **G and H)** Low (G) and high (H) magnification stereo fluorescence microscopy imaging of red fluorescent protein (TdTomato) expressed in the arterial vasculature of the heart.

The preservation of *in vivo* dyes was next assayed by performing intravital dye labeling of blood conducting vessels in neonatal murine pups^35,36^, when complex vascular remodeling is occurring, notably in the eye. Half-mounted heads were isolated, optically cleared with ATC-OC, and imaged by confocal microscopy. Results showed a robust preservation of fluorescently labeled cranial and facial vessels throughout the head (Fig. 11A), and high magnification analyses provided a detailed 3D view of the intricate vascular bed perfusing the neonatal eye cup and retina (Fig. 11B and Supplemental Movie 7). ATC-OC was also proven to provide a sophisticated modality to characterize an organ’s cellular structures when immunofluorescence was combined with intravital dye labeling. This is shown in Fig. 11C and D, where the vasculature of exposed neonatal torsos was immunolabeled post intravital dye labeling. The intravital dye labeling (red) was readily distinguishable from the immunolabeled vasculature (green, Endomucin), and at high magnification enabled the visualization of mature vessels that conduct blood (yellow) versus immature vessels (green only) that are not yet perfused (Fig. 11D, and Supplemental Movie 8).

**Figure 11.**
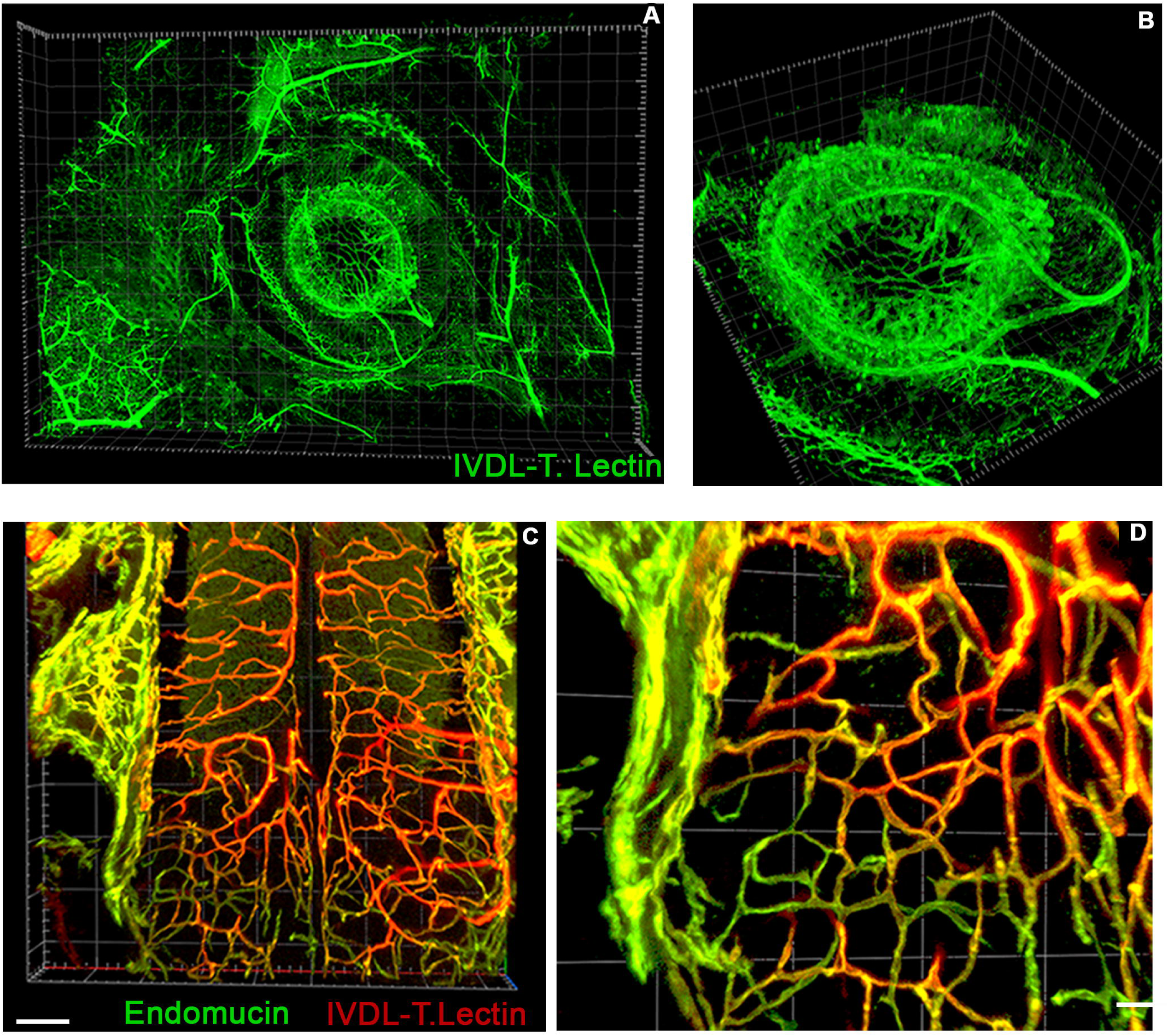
Imaging of *in vivo* tracers in conjunction with IF using ATC. **A**, **B)** 3D imaging of half mounted neonatal murine heads cleared with ATC-OC after intravital dye labeling (IVDL, tomato lectin) of the circulatory system validated compatibility with *in vivo* tracers (A, facial vasculature; B, high magnification of the eye vasculature; also see Supplemental Movie 7). **C, D)** Confocal imaging of neonatal body trunks cleared with ATC-OC after intravital dye labeling of blood conducting vessels (red) and IF for the whole vasculature (vascular endothelium, endomucin, green). High magnification analyses (D) enabled visualization of patent blood conducting vessels (yellow) and nascent unperfused vessels (green; also see Supplemental Movie 8).

ATC-OC was further used to interrogate human kidney biopsies (Fig. 12). Due to its more robust proteinaceous extracellular and cellular consistency, human tissue is well known to be tougher and more autofluorescent than counterpart tissue from smaller mammalian animal models that are commonly used for research, such as mice^1^. To this end, we aimed to determine if the different tissue layers of the glomerulus could be visualized in human samples, as was possible with murine renal tissue. Human samples were stained for Podocalycin, PDGFR-β, and CD31 to mark the podocyte, mesangium and endothelial layers of the glomerulus, respectively. All three tissue layers were distinctly imaged, with single cell resolution (Fig. 12A-E). Thus, ATC-OC served to 3D image both murine and human tissue samples at high resolution.

**Figure 12.**
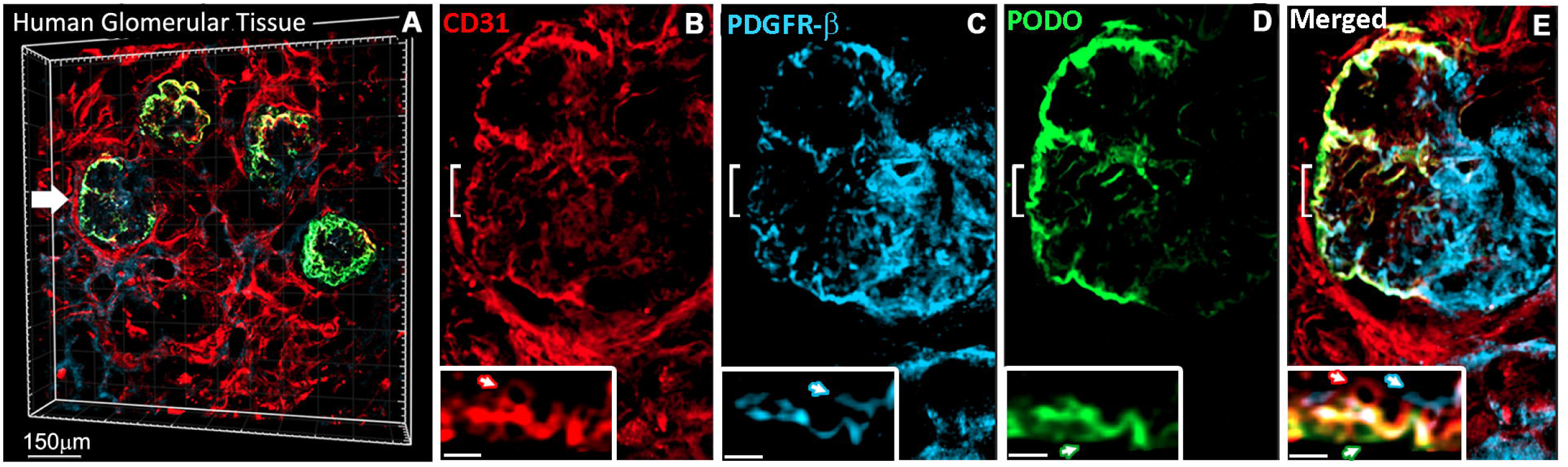
ATC facilitates multiplex IF imaging of human renal tissue. ATC was used to perform 3D multiplex imaging of human tissues that exhibit notoriously high AF. **A-E)** Human samples immunolabeled for the glomerular tissue layers (A, merged labeled image, arrow points to region shown at high magnification), including the endothelial (B, CD31, red), mesangial (C, PDGFR-β, blue), and the podocyte (D, podocalyxin) tissue regions (E, high magnification merged image). Notably, all tissue regions could be distinctly visualized, with cellular resolution (inset shows high magnification of region demarked by white brackets in panels F-I).

## Discussion

We developed ATC, an aqueous optical clearing platform designed specifically to address limitations associated with 3D imaging of tissues that naturally exhibit high AF, including those such as the heart that are both highly AF and are difficult to clear. Our studies demonstrate AT-HP provides advantageous properties that include superior imaging resolution with SNRs that are up to 200% that of leading 3D imaging methods; increased fluorescent reporter protein signal detection; compatibility with up to 10-fold lower IF antibody concentrations; and a significant increase in the speed and efficacy of refractive index matching solutions to render tissue transparent (nine total tested). As we show, these properties make ATC-OC ideal for performing imaging studies of challenging tissue types such as the kidney, testis, and heart that exhibit robust AF. ATC-OC facilitated studies such as visualization of the previously undocumented architecture of adjacent connecting and collecting tubule nephron segments, utilization of AI based auto-segmentation workflows with simple, low tech microscopy, detection of poorly recognized adult spermatogonial stem cell markers, and the anatomic localization of the sinoatrial pacemaker node of the heart. Cumulatively, our findings establish ATC-OC as a powerful tool for researchers to perform complex 3D imaging studies that, to date, have remained challenging.

To optimize the imaging of autofluorescent and fibrous tissues, we designed ATC-OC as a two-stage methodology, with an initial tissue treatment step using AT-HP that serves to decrease tissue AF. We demonstrate that AT-HP treatment significantly decreases AF in a time dependent manner. In heart tissue, for instance, green field AF was quenched by up to 60% and red AF nearly abolished within 48 hr. Notably, the time dependent quenching serves as a useful property of AT-HP, as investigators can vary the time of use to fit the AF of specific samples. This will be particularly useful for studies interrogating wide-ranging samples, including those assaying human tissues that are known to exhibit especially high background. As shown, we leveraged AT-HP to image both murine and human kidney samples with advanced resolution, including markers that are not readily detected using current methods.

We established AT-HP also significantly augments optical clearing. Live imaging showed near instantaneous optical clearing of 0.5mm heart and kidney sections using AT-HP in combination with EZ View, CUBIC-2, MACS-R2 and FOCM. Untreated tissues took longer to begin clearing, and never reached the same level of transparency. Indeed, we corroborated a significant increase in clearing potency with all of the refractive index matching solution we tested, nine in total. Strikingly, AT-HP followed by EZ View was able to readily clear whole mouse hearts and kidneys in as fast as three hours. By contrast, EZ Clear reported requiring 24 hours of refractive index matching for hearts and kidneys with modest transparency, thus marking a 8x increase in speed of clearing using ATC-EZ View. Notably, these data demonstrate that chemical delipidation, for example with THF, is not requisite for fast and efficacious optical clearing. This further adds to the benefits of AT-HP, as it can be integrated into already established working protocols to both mitigate AF and enhance tissue transparency, as with ATC-EZ View.

In addition to AT-HP, we also developed AT-OC, a refractive index matching solution comprised of formamide and glycerol. Formamide was chosen for optical clearing as we discovered it produced a moderate reduction in AF. By contrast, similar benefits were not observed when using comparative 3D imaging technologies (see supplementary Fig. 2 and 5). We demonstrate that the combination of glycerol with formamide provides greater tissue clearing than protocols such as ClearT^2^ that also use formamide. Moreover, the sequential use of AT-HP and then AT-OC, or ATC-OC, provides sufficient optical clearing with high preservation of tissue morphology, serving to render transparent whole murine muscular organs, including the heart, stomach, upper urinary system, intestine, and bladder. It should be noted that, aside from the benefits of ATC-OC, it was not as fast or potent in rendering tissue transparent, as say ATC-EZ View.

The mitigation of tissue background noise, as is seen with AT-HP or AT-OC, is only beneficial if it is consistent with an increase in SNRs. To this end, we show AT-HP provides significantly higher SNRs than standard IF, and remarkably facilitates the use of AI auto-segmentation workflow with low tech microscopy technology such as stereo fluorescence microscopy (e.g. no optical sectioning). Moreover, we demonstrate AT-HP provides significantly higher SNRs than CUBIC, EZ Clear, iDISCO+, ScaleS, and Passive Clarity (PACT). SNRs were shown to be as much as 200% of SNR obtained with these methodologies. Consistent with these findings, the high resolution and SNRs provided by ATC-OC facilitated complex 3D imaging studies. ATC-OC permitted multiplex imaging of renal nephrons that are among the most autofluorescent tissue structures, enabling the visualization of adjacent connecting and collecting tubule segments with single cell resolution. In the testis, ATC-OC enabled visualization of cKit, Stra8, and GFRα1, spermatogonial stem cell markers that are shown to be poorly recognized by IF. This allowed the visualization of varying stages of stem cell differentiation in whole testis tubules. ATC-OC also permitted imaging 1.6 mm into the heart by confocal microscopy, and allowed visualization of the HCN4+ sinoatrial pacemaker node that was not readily observed by comparative methods. Finally, we demonstrate that the high SNRs obtained with ATC-OC enable an up to 10-fold lower primary antibody concentration, as shown in IF studies of the heart, kidney and testis.

Remarkably, to the best of our knowledge, ATC is the first 3D imaging technique that improves the detection of fluorescent reporter proteins. AT-HP significantly resolved the signal from RFP+ arteries in transgenic heart samples, with SNRs increasing by up to 64%. These findings were corroborated in transgenic kidneys showing Venus+ interstitial cells exhibit >50% higher SNRs. Notably, in the testis, testicular Venus+ interstitial cells that were not discernible from background tissues due to their low Venus expression became readily detectable using AT-HP based IF, and SNRs were found to be a striking 5.8x higher. These finding were consistent across model systems, as shown in ATC-OC imaging of GFP-expressing stem cell-derived EBs and GFP-expressing zebrafish hearts. Moreover, ATC-OC was shown to be compatible with fluorescent tracers. Using intravital dye labeling, we were able to visualize the complex 3D architecture of the ocular vasculature, as well as nascent and patent vessels of the murine trunk. Thus, ATC-OC can serve as a valuable tool for the large community of researchers employing fluorescent tracers and wide-ranging transgenic animal models that express fluorescent report proteins.

In summary, we developed ATC-OC, a 3D imaging technology that provides increased imaging resolution and SNRs, greater detection of poorly recognized antigens, augmented optical clearing, and enhanced detection of fluorescent reporters. These properties facilitate 3D interrogation of challenging tissue types, and thus make ATC a valuable new tool for performing complex imaging studies.

## METHODS

### Animals

Adult mice were purchased from Taconic Farms (Germantown, NY, USA). Spry4^H2B-Venus^ mice expressing Venus fluorescent reporter protein from the endogenous Spry4 locus has been previously described^19^. The transgenic zebrafish Tg(*myl7:EGFP*) reporter line has been described^36^. All animals were housed at an AAALAC accredited facility at the Weill Medical College of Cornell University Animal Facility. All experiments were approved and carried out in accordance with the ethical guidelines and regulations stipulated by the Institutional Animal Care and Use Committee of Weill Medical College of Cornell University.

### Atacama Clear

Organs were immersion fixed with 4% paraformaldehyde (PFA) at 4°C using a rotisserie-style shaker. Fixation time was modulated according to tissue thickness, and whether organs were of soft tissue or muscular tissue type. Sufficient time of fixing was required for adequate preservation of antigens. Three week-old whole adult mouse hearts and kidneys were fixed for 72 hr, mouse neonatal tissues and adult zebrafish hearts were fixed for 24 hr, and stem cell-derived EBs were fixed for 3 hr. After fixation, it was critical to wash free fixative to obtain reproducible immunostaining results. To this end, tissues were rinsed with copious amounts of PBS, then washed at 4 ^O^C for at least 48 hr with PBS, and then stored in PBS until use. Note, all steps of fixing and ATC protocol are done using rotisserie-style shaker to ensure proper penetration of solutions.

ATC consists of two stages; stage 1 is treatment with AT-HP, followed by stage 2, or treatment with AT-OC. The base solution of AT-HP consists of the following final concentrations: 150mM KCl (w/v), 0.5% Triton-X (v/v), 20% DMSO (v/v), 0.5% NaN3 (w/v), 10 mM Tris-HCl (w/v), pH 9.5 made in basic double distilled H_2_0 (pH 9 with NaOH; final pH of base solution is 9.5). Base solution is mixed well, and then allowed to settle for 10 minutes. Right before use, H_2_O_2_ is directly added to AT-HP base solution. To start, a final concentration of 0.05% H_2_O_2_ is always used (diluted from 30% stock solution v/v, Sigma, H1009). AT-HP containing 0.05% H_2_O_2_ is mixed well, and the allowed to acclimate at RT for 10 minutes. Organs are then submerged in AT-HP within a suitable receptacle, such as an Eppendorf tube or cryogenic vial, and gently rocked on a rotisserie shaker at room temperature. Tissues are observed to ensure that air pockets are not accumulating within the tissue; minor bubbling at the edges of tissues is permissible. In instances where robust oxygenation was observed, the reaction was stopped by rinsing tissue in PBS, and the reaction mixture was adjusted to proceed more slowly by lowering the concentration of H_2_O_2_ to 0.01%. Once oxygenation was confirmed to be lacking, samples were incubated in AT-HP at room temperature for up to 4 hr. Tissues were then rinsed 2x and incubated with wash buffer comprised of 150 mM KCl, 10% DMSO, 0.5% triton-x, and 20 mM Tris HCl, pH 9.5 dissolved in basic DDH_2_0 for 20 min – 1 hr at 37 ^O^C. Tissue decolorization and autofluorescence was then assessed. If further processing is needed, a new round of fresh AT-HP treatment was done by increasing the concentration of H_2_O_2_ by 0.05%. In this manner, AT-HP treatment can continue until desired results by increasing the concentration of H_2_O_2_. The maximum concentration of H_2_O_2_ required for our studies was 1% (all animal tissue were treated at a maximum concentration of 0.5% H_2_O_2_; human samples were treated with a max concentration of 1% H_2_O_2_). In between AT-HP treatments, samples are left rocking in wash buffer at room temperature (for example overnight, so as to continue AT-HP treatment on day 2). Optionally, to expedite quenching and treat samples marked by particularly high AF, tissues can be incubated at room temperature overnight with a priming solution consisting of 1% glycerol (v/v), 0.5% Triton-x (v/v), and 20mM Tris-HCl (w/v) in basic H_2_O prior to initial treatment of AT-HP. Finally, although all the data in the present study were obtained with room temperature AT-HP treatment, we did observe in test studies that AT-HP quenching of AF could be maximized by going from room temperature to 37 ^O^C, starting back again at 0.05%. However, quenching should be monitored more closely at 37 ^O^C and should be avoided with tissues expressing endogenous fluorescent reporters.

After AT-HP treatment, samples were amenable to previously established standard immuno-staining protocols. For our studies, tissues were rinsed 3 x 5 min with PBS + 0.5% triton-x (PBST), and then permeabilized in blocking solution consisting of PBST + 1% NDS + 1% DMSO overnight at 37 ^O^C. Primary antibodies were then incubated in PBST + 1% DMSO. Antibodies used in this study were: anti-α-smooth muscle actin (F3777 clone 1A4, Sigma, for titration studies standard concentration of 1:200 of 1mg/ml stock concentration; for comparative studies of 3D imaging protocols, at 1:1000), anti-ENAC (Ergonul et al., at 1:200)^39^, anti-parvalbumin (R&D AF5058, at 1:200), anti-endomucin (ab106100, Abcam, at 1:200), anti-podocalyxin (3169, R&D Systems, at 1:200), anti-HCN4 (APC-052, Alomone, at 1:500), anti-Pdgfr-β (3169, Cell Signaling Technology, at 1:100), anti-CD31 (BD Biosciences, 550389 at 1/500), anti-E-cadeherin (AF748, R&D, for titration studies standard concentration of 1:200 of .2mg/ml stock), anti-MCAM (134702, BioLegend, for titration studies standard concentration of 1:500 of .5mg/ml stock), and cardiac Troponin-T (NB110-2546, R&D Systems, at 1:500). Minimally cross-absorbed Alexa Flour secondary antibodies were used to detect primary antibodies.

Optical clearing with AT-OC was next performed to render immunostained tissue transparent. Optical clearing with AT-OC consisted of impregnating samples with sequential solutions of varying formamide/glycerol concentrations, with each solution rendering tissue increasingly more transparent (deionized formamide, Sigma, F7503; ultrapure glycerol, ThermoFisher, AC410985000). Specifically, tissues were immersed with the following the following sequence of solutions made in basic DDH_2_0 at room temperature: 20% formamide + 0.5% Triton-x, 20% formamide, 20% formamide/5% glycerol, 20% formamide/10% glycerol, 20% formamide/25% glycerol, 20% formamide/ 50% glycerol, 30% formamide/70% glycerol, and then a final solution comprised of 17% formamide + 5% Triethalomine + 70% glycerol + 3% Tris base (w/v, Tris, Promega, H5135) made in basic DDH_2_0. At the onset, tissues are incubated with 20% Formamide + .5% Triton-x overnight at room temperature to prime the tissue for optical clearing. Traces of Triton-x are then removed by washing with 20% Formamide 3x. The time of incubation in each subsequent solution varied depending on the tissue being cleared, in particular because the viscosity of clearing solutions increases with rises in glycerol concentration. Thus, care is needed to ensure that the entire tissue has been impregnated with the current solution before moving forward to subsequent solutions. Penetration of clearing mixtures into samples could be ascertained by ensuring samples sank to the bottom of containers. For large organs, dabbing tissue clear of solutions before adding the subsequent mixture further facilitated optical clearing. As a guideline, the number of solutions could be reduced to enable faster clearing if working with thinner or sectioned tissue. In such instances, treatment could start with 20% formamide +.5% triton-x, then 20% formamide alone, followed by quarter percent increases of glycerol, starting with 20% formamide/25% glycerol. Moreover, the entire sequence of solutions was not needed to optically clear all tissue, e.g. 0.5 mm kidney sections were rendered transparent with 20% formamide/25% glycerol. After clearing, tissues were imaged directly in final clearing solution, using glass bottom plates with spacers, and then placing a coverslip over the sample to ensure it does not move. Samples could be stored in final clearing solution for months. Confocal microscopy was performed on a Zeiss LSM 800 and stereo fluorescence microscopy on a Nikon SMZ1500 mounted with a Nikon DS-FI3 camera. 3D and AI analyses were performed using Imaris Imaging Software (Bitplane). For quantification of transparency, tissue clarity was quantified using the gradient magnitude of the background grid as a measure of how well the grid showed through the tissue. Gradient magnitude of the image was calculated in Matlab using a Prewitt operator. Grid visibility ratios were computed by assaying horizontal edges in each image and dividing the gradient magnitude of the grid edge behind the tissue section by the gradient magnitude of the grid edge adjacent to the tissue. A ratio of one indicates perfect transparency and a ratio of zero indicates complete opacity.

### Intravital Dye Labeling

Intravital dye labeling was performed as previously described^38^. In short, 100 µl of fluorescently tagged tomato lectin (0.2mg/ml in PBS, Vector laboratories) was injected into the circulation of anesthetized newborn pups (postnatal day zero) by intracardiac injection. Pups were euthanized after a 5 min period to let the lectin circulate, and tissues harvested and fixed.

### Stem cell-derived embryoid bodies

Mouse embryonic stem cells (mESCs) with a doxycycline inducible GFP transgene^40^ were maintained on gelatin coated tissue culture treated plates in serum-free conditions. Media (2i plus LIF) was composed of 50% Neurobasal media (Gibco) and 50% DMEM/F12 (Corning cellgro) supplemented with 0.5X N2 and 0.5X B27 (Gibco), 100U/mL Penicillin, 0.1 mg/ml Streptomycin (Corning), 0.05% BSA (Gibco), 2 mM Glutamine (Corning), 0.15 mM monothioglycerol (Sigma), 1000U/ml mouse LIF (ESGRO, Sigma), 3 uM CHIR99021 (Stem Cell Technologies), and 1uM PD0325901 (Stem Cell Technologies). To produce embryoid bodies (EBs) mESCs were differentiated in serum free differentiation media (SFD). SFD consisted of 75% IMDM (Gibco), 25% Ham’s F12 (Corning cellgro), 0.5X N2 (Gibco), 0.5X B27 (Gibco), 0.05% BSA (Gibco), 0.5mM Ascorbic Acid (Sigma), 2mM Glutamine (Corning), 0.45 mM monothioglycerol (Sigma), 100U/mL Penicillin and 0.1 mg/ml Streptomycin (Corning). On the first day of differentiation, mESCs were dissociated with Accutase (Sigma) and plated at 40,000 cells/ml in 12.5ml of SFD in 100mm petri dishes. After 48 hr, EBs were dissociated with Accutase and plated again at 80,000 cells/ml in SFD supplemented with 75ng/ml Activin A (R&D Systems). After 24 hours, expression of GFP was induced by adding Doxycycline (Sigma) for 24 hours before EBs collection.

## Supporting information

Live Imaging of Optical Clearing with ATC-EZ View

Live Imaging of Optical Clearing with ATC-CUBIC-2

Live Imaging of Optical Clearing with ATC-MACS-R2

Live Imaging of Optical Clearing with ATC-FOCM

3D Imaging of Glomerular Tissue Layers

3D Imaging of Renal Nephrogenic Zones

3D Imaging of Facial Vasculature

3D Imaging of Perfused and Nascent Vasculature of the Trunk

## ACKNOWLEDGEMENTS

This work was supported by an NIH NIDDK R21DK116171, R01DK133381 grant and Mastercard Diversity-Mentorship Collaborative Pilot Grant to R.H. Support to N.D.S. was by the Weill Cornell Hartman Foundation for Therapeutic Organ Regeneration. We thank Dr. Miriam Gordillo for providing EB samples. We thank Dr. Lawrence Palmer for providing advice on nephron staining and ENAC antibody.

## AUTHOR CONTRIBUTIONS

Study conception, R.H.; Technological Conception and Development, R.H.; Experiment Design, R.H., T.E., E.A.J, L.L.; Experiments, R.H., T.C., L.L, N.D.S., and Y.L.; Intellectual Contributions, R.H., T.E., E.A.J.; Manuscript Writing, R.H. with input from T.E., E.A.J, and L.L.

## DECLARATION OF INTERESTS

R.H. is inventor on a U.S. patent relating to optical clearing technology submitted by the Center for Technology Licensing at Cornell University.

## FIGURE LEGENDS

**Supplementary Figure 1.**
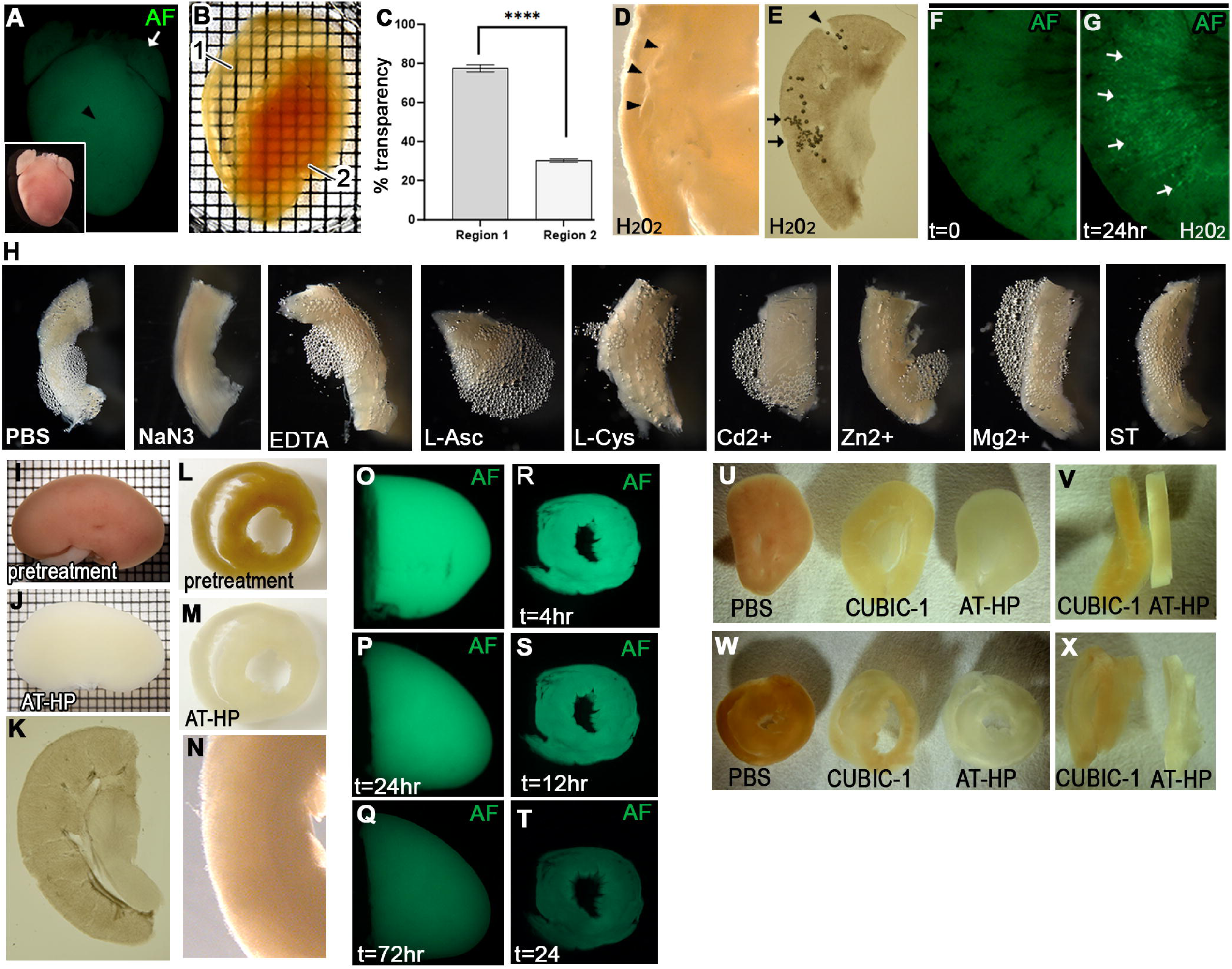
Development of AT-HP, a robust method for quenching tissue AF while preserving tissue morphology. **A)** Bright field image (inset) and green AF of perfused hearts showed that regions of the organ that retained robust amounts of blood, such as the ventricle (black arrowhead), exhibited higher AF than regions that were readily cleared of blood, such as the atria (white arrow). **B)** Optically cleared heart showing regions devoid of blood are rendered significantly more transparent (region 1) than regions flush with blood that are resistant to clearing (region 2), as illustrated by the visibility of the underlying grid below the heart. **C)** Quantification of transparency in regions 1 and 2 of cleared heart (p<0.0001). **D, E)** Bright field images of heart (D) and kidney (E) tissue morphology after treatment with standard H_2_O_2_ (arrow heads point to sites of tissue tearing; arrows point to oxygen pockets trapped within tissue). **F, G)** AF of kidney tissue before (F) and after (G) standard H_2_O_2_ treatment. Arrows point to regions of tissue damage after H_2_O_2_ treatment that then exhibited increased AF. **H)** Screen for inhibitors of H_2_O_2_ generated tissue oxygenation; heart sections treated with H_2_O_2_ plus sodium azide (NaN3), EDTA, L-ascorbic acid (L-Asc), L-cysteine (L-Cys), cadmium (Cd^2+^), zinc (Zn^2+^), magnesium (Mg^2+^), or sodium thiosulfate (ST). **I-K**) Images of a whole kidney before (I) and after (J) treatment with AT-HP. (K) Bright field image of representative kidney section morphology after treatment. **L-N)** Images of a 1 mm heart sections before (L) and after (M) treatment with AT-HP. (N) Bright field image of representative heart section morphology after treatment. **O-Q)** Time dependent decrease in AF of halved kidneys treated with AT-HP. **R-T)** Time dependent decrease in AF of 1 mm heart sections treated with AT-HP. **U-X)** A single kidney (U and V) and heart (W and X) were cut into 0.5 mm sections, and adjacent sections treated with AT-HP or CUBIC-1. Bright field images of adjacent sections side-by-side in the same field of view were taken for comparison of tissue morphology post decolorization.

**Supplementary Figure 2.**
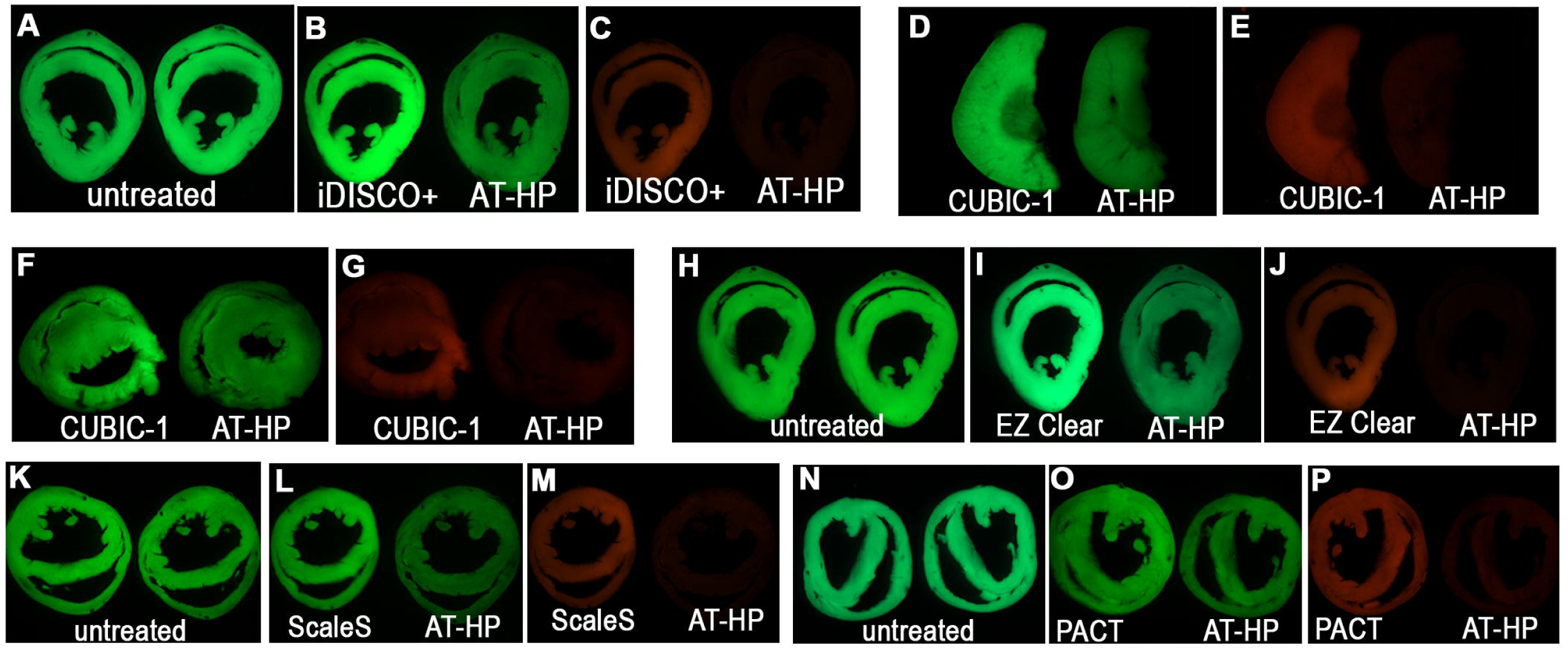
AF of tissues after treatment with AT-HP or widely used optical clearing protocols. 0.5 mm thick sections obtained from a single heart or kidney were treated with AT-HP or comparative 3D imaging protocol and green and red AF assessed prior to beginning IF labeling, including iDISCO+ (A-C), CUBIC (D and E, kidney; F and G, heart), EZ Clear (H-J), Sca*l*eS (K-M), and PACT (N-P).

**Supplemental Figure 3.**
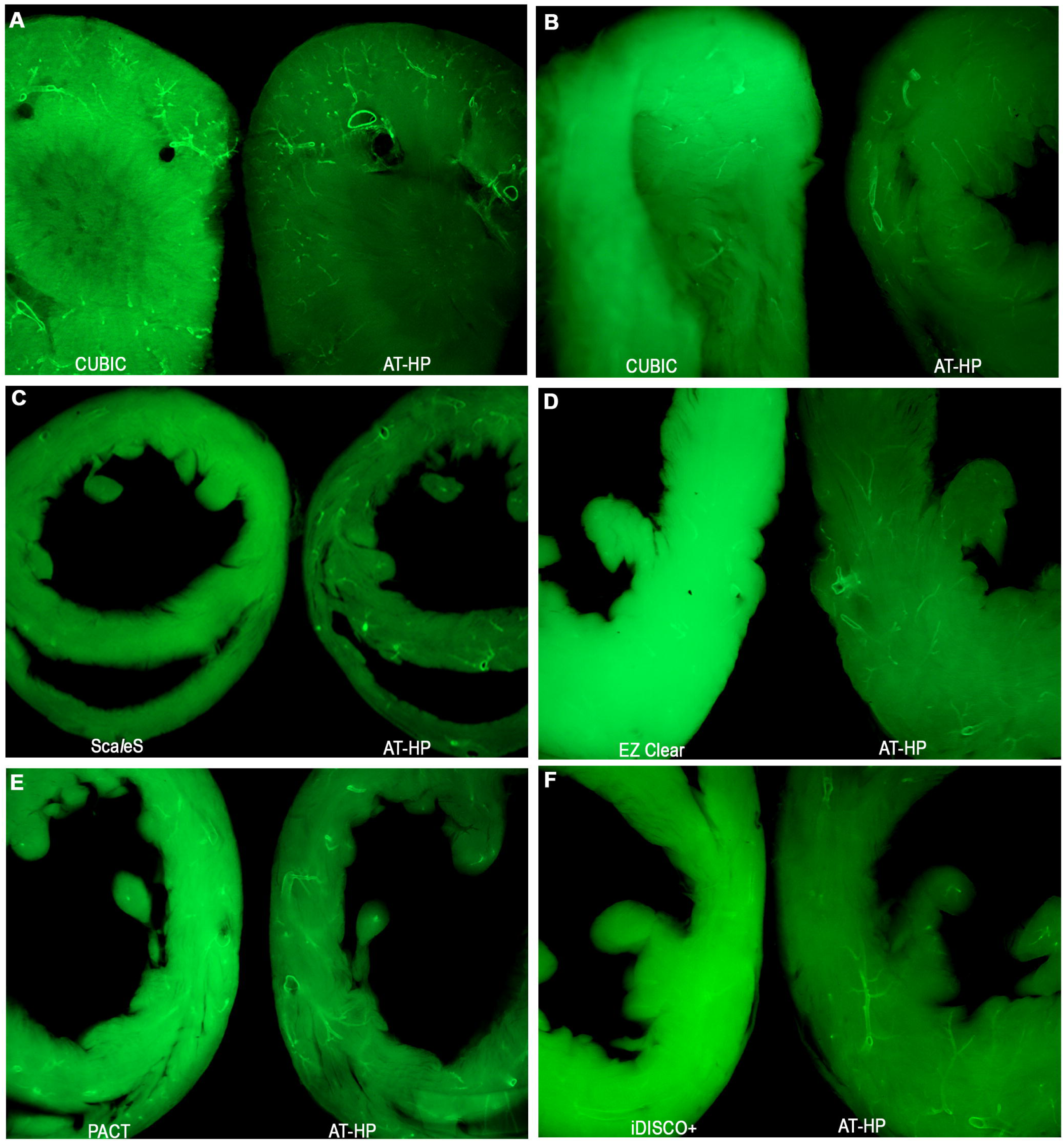
AT-HP provides superior IF image resolution than widely used 3D clearing methods. **A-F)** Individual kidneys (A) or hearts (B-F) were cut into 0.5 mm thick sections. IF was then performed on adjacent sections for the arterial vasculature (SMA, green) using AT-HP or CUBIC (A and B), Sca*l*eS (C), EZ Clear (D), PACT (E), or iDISCO+ (F). Stereo fluorescence microscopy images of immunolabeled sections were taken side-by-side, in the same field of view, for direct comparison.

**Supplemental Figure 4.**
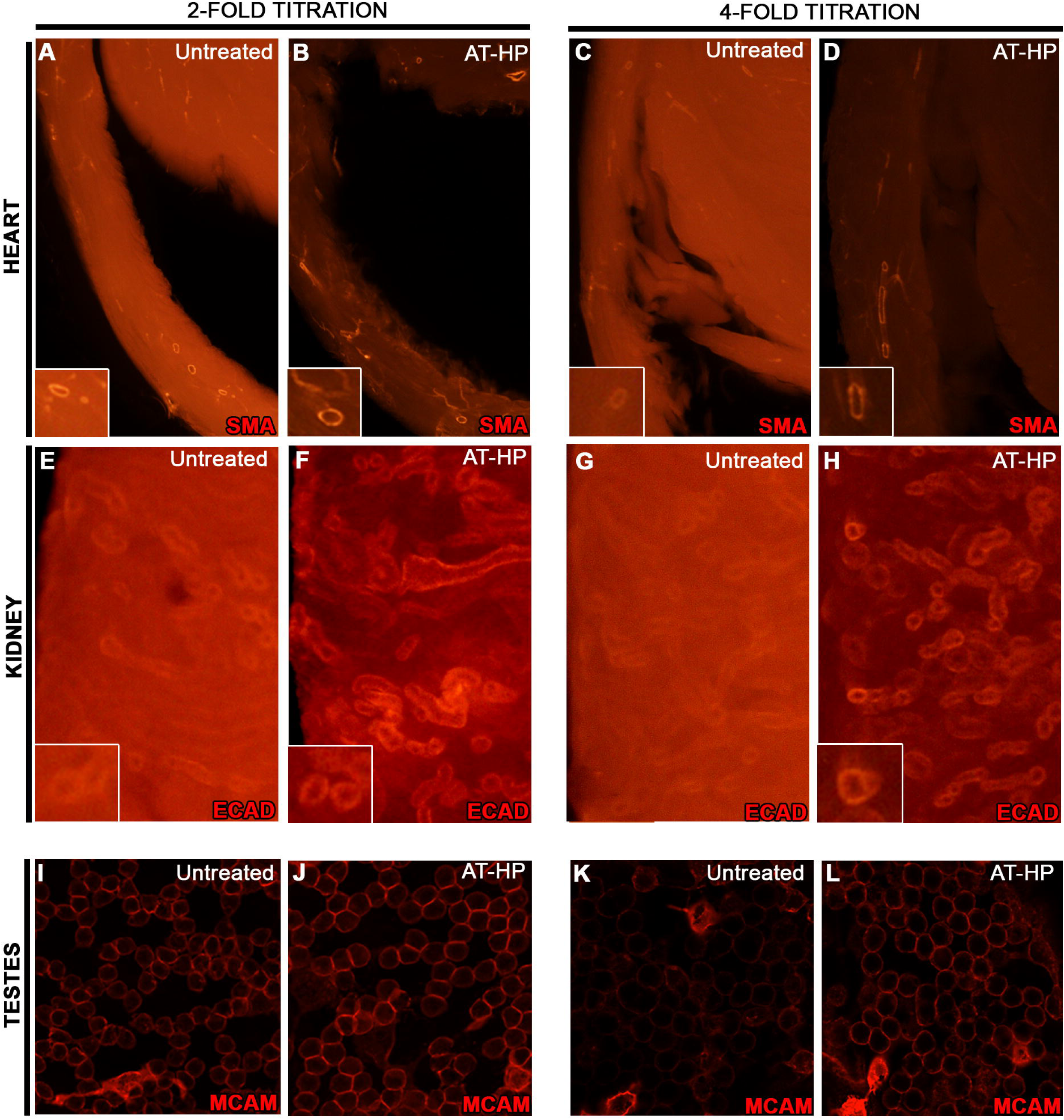
AT-HP facilitates the use of lower antibody concentrations in IF. IF using highly published antibodies on heart, kidney and testis tissues. **A-D)** IF for the arterial vasculature (SMA, red) on a halved heart section, where one half was left untreated and the second was treated with AT-HP, using two-fold lower (A and B) or 4-fold lower antibody concentrations (C and D) than standard published concentrations (see Fig. 3). **E-H)** IF for the renal collecting tubules (ECAD, red) on a halved kidney section, where one half was left untreated and the other treated with AT-HP, using two-fold lower (E and F) or 4-fold lower antibody concentrations (G and H) than standard published concentrations. **I-L)** IF for germ cells (MCAM, red) on testis sections that were left untreated or treated with AT-HP using two-fold lower (I and J) or 4-fold lower antibody concentrations (K and L) than standard published concentrations.

**Supplemental Figure 5.**
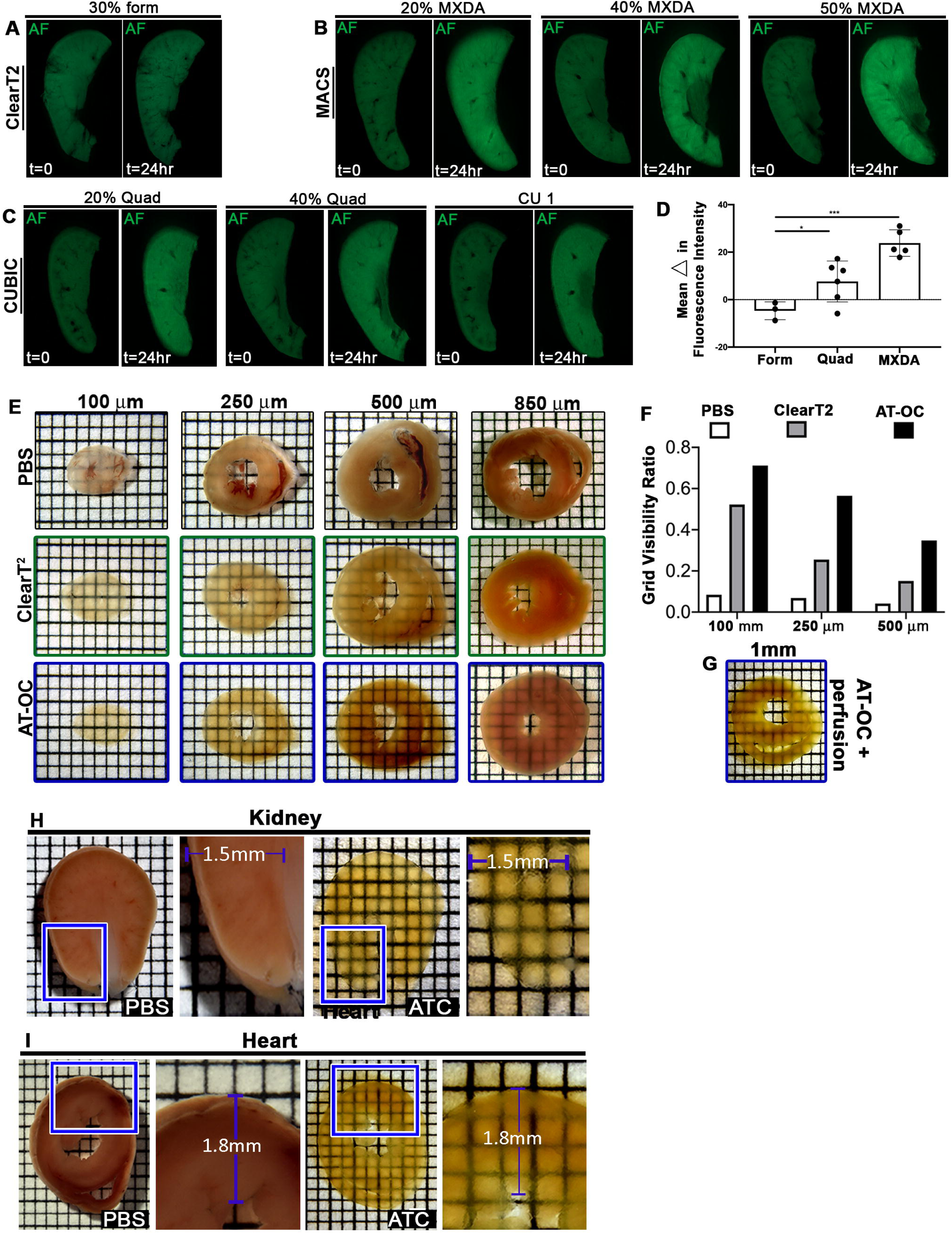
Development of ATC-OC. **A-C)** Green AF of tissue before and after treatment with reagents from ClearT^2^ (A, formamide), MACS (B, m-Xylylenediamine, MXDA), and CUBIC (C, quadrol) optical clearing techniques. **D)** Quantification of mean change in tissue AF after treatment with optical clearing reagents. **E)** Optical clearing of heart sections of different thickness with ClearT^2^ or AT-OC. **F)** Quantification of optical clearing with ClearT^2^ or AT-OC. **G)** Optical clearing of 1 mm thick heart section after aortic arch perfusion, and then treatment with AT-OC. **H, I)** Measurements of 1mm thick kidney (H) and heart (I) sections before and after treatment with ATC showed high tissue preservation of morphology and size.

**Supplementary Figure 6.**
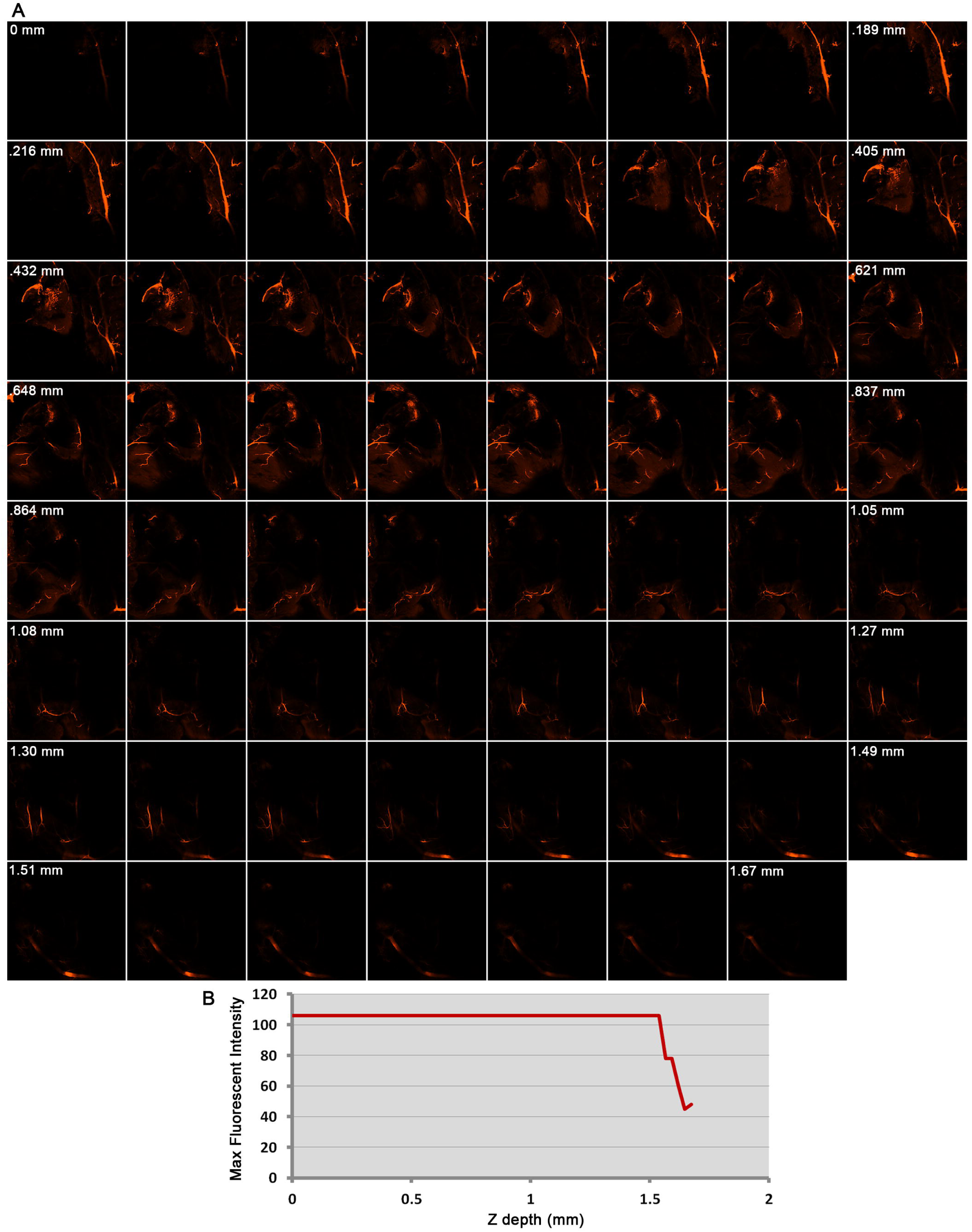
ATC preserves IF signal intensity when interrogating deep inside organs. A) All representative optical slices of a 3D heart image (see Fig. 8 for 3D reconstruction) performed using ATC-OC. B) Quantification of signal intensity shows IF signal was preserved without loss up to the confocal optical working distance of 1.6mm.

**Supplemental Movie 1:** Live imaging of optical clearing with ATC-EZ View

**Supplemental Movie 2:** Live imaging of optical clearing with ATC-CUBIC-2

**Supplemental Movie 3:** Live imaging of optical clearing with ATC-MACS-R2

**Supplemental Movie 4:** Live imaging of optical clearing with ATC-FOCM

**Supplemental Movie 5:** 3D image of glomerular tissue

**Supplemental Movie 6:** 3D image of nephrogenic tubule zones

**Supplemental Movie 7:** 3D image of facial vasculature

**Supplemental Movie 8:** 3D image of perfused and nascent vasculature of the trunk

## Notes

### Summary of Updates

This version of the manuscript has been revised with data demonstrating Atacama Clear: a) facilitates use of AI auto-segmentation with simple stereo microscopy; b) increases 3D imaging SNRs of endogenous fluorescent reporters; c) potentiates the optical clear capacity of refractive index matching solutions (increased benchmarking); d) facilitates the use of up to 10x less antibody concentration; and e) significantly decreases the time of optical clearing.

## REFERENCES

1 Richardson, D. S. & Lichtman, J. W. Clarifying Tissue Clearing. Cell 162, 246–257, doi:10.1016/j.cell.2015.06.067 (2015).

2 Yu, T., Zhu, J., Li, D. & Zhu, D. Physical and chemical mechanisms of tissue optical clearing. iScience 24, 102178, doi:10.1016/j.isci.2021.102178 (2021).

3 Hama, H. et al. Scale: a chemical approach for fluorescence imaging and reconstruction of transparent mouse brain. Nat Neurosci 14, 1481–1488, doi:10.1038/nn.2928 (2011).

4 Hama, H. et al. ScaleS: an optical clearing palette for biological imaging. Nat Neurosci 18, 1518–1529, doi:10.1038/nn.4107 (2015).

5 Kuwajima, T. et al. ClearT: a detergent– and solvent-free clearing method for neuronal and non-neuronal tissue. Development 140, 1364–1368, doi:10.1242/dev.091844 (2013).

6 Ke, M. T., Fujimoto, S. & Imai, T. SeeDB: a simple and morphology-preserving optical clearing agent for neuronal circuit reconstruction. Nat Neurosci 16, 1154–1161, doi:10.1038/nn.3447 (2013).

7 Hou, B. et al. Scalable and DiI-compatible optical clearance of the mammalian brain. Front Neuroanat 9, 19, doi:10.3389/fnana.2015.00019 (2015).

8 Susaki, E. A. et al. Whole-brain imaging with single-cell resolution using chemical cocktails and computational analysis. Cell 157, 726–739, doi:10.1016/j.cell.2014.03.042 (2014).

9 Tainaka, K. et al. Whole-body imaging with single-cell resolution by tissue decolorization. Cell 159, 911–924, doi:10.1016/j.cell.2014.10.034 (2014).

10 Matsumoto, K. et al. Advanced CUBIC tissue clearing for whole-organ cell profiling. Nature protocols 14, 3506–3537, doi:10.1038/s41596-019-0240-9 (2019).

11 Renier, N. et al. Mapping of Brain Activity by Automated Volume Analysis of Immediate Early Genes. Cell 165, 1789–1802, doi:10.1016/j.cell.2016.05.007 (2016).

12 Hsu, C. W. et al. EZ Clear for simple, rapid, and robust mouse whole organ clearing. Elife 11, doi:10.7554/eLife.77419 (2022).

13 Azaripour, A. et al. A survey of clearing techniques for 3D imaging of tissues with special reference to connective tissue. Progress in histochemistry and cytochemistry 51, 9–23, doi:10.1016/j.proghi.2016.04.001 (2016).

14 Akiyama, F. et al. A multiwell plate approach to increase the sample throughput during tissue clearing. Nature protocols, doi:10.1038/s41596-024-01080-1 (2024).

15 Renier, N. et al. iDISCO: a simple, rapid method to immunolabel large tissue samples for volume imaging. Cell 159, 896–910, doi:10.1016/j.cell.2014.10.010 (2014).

16 Murphy, C. & Lazzara, R. Current concepts of anatomy and electrophysiology of the sinus node. Journal of interventional cardiac electrophysiology: an international journal of arrhythmias and pacing 46, 9–18, doi:10.1007/s10840-016-0137-2 (2016).

17 Sartiani, L., Mannaioni, G., Masi, A., Novella Romanelli, M. & Cerbai, E. The Hyperpolarization-Activated Cyclic Nucleotide-Gated Channels: from Biophysics to Pharmacology of a Unique Family of Ion Channels. Pharmacol Rev 69, 354–395, doi:10.1124/pr.117.014035 (2017).

18 Ludwig, A. et al. Two pacemaker channels from human heart with profoundly different activation kinetics. The EMBO journal 18, 2323–2329 (1999).

19 Morgani, S. M. et al. A Sprouty4 reporter to monitor FGF/ERK signaling activity in ESCs and mice. Dev Biol 441, 104–126, doi:10.1016/j.ydbio.2018.06.017 (2018).

20 Luo, Y. et al. SPRY4-dependent ERK negative feedback demarcates functional adult stem cells in the male mouse germlinedagger. Biology of reproduction 109, 533–551, doi:10.1093/biolre/ioad089 (2023).

21 Zhu, X. et al. Ultrafast optical clearing method for three-dimensional imaging with cellular resolution. Proc Natl Acad Sci U S A 116, 11480–11489, doi:10.1073/pnas.1819583116 (2019).

22 Yu, T. et al. RTF: a rapid and versatile tissue optical clearing method. Scientific reports 8, 1964, doi:10.1038/s41598-018-20306-3 (2018).

23 Chung, K. et al. Structural and molecular interrogation of intact biological systems. Nature 497, 332–337, doi:10.1038/nature12107 (2013).

24 Marx, V. Microscopy: seeing through tissue. Nature methods 11, 1209–1214, doi:10.1038/nmeth.3181 (2014).

25 Pollak, M. R., Quaggin, S. E., Hoenig, M. P. & Dworkin, L. D. The glomerulus: the sphere of influence. Clin J Am Soc Nephrol 9, 1461–1469, doi:10.2215/CJN.09400913 (2014).

26 Nagata, M. Glomerulogenesis and the role of endothelium. Curr Opin Nephrol Hypertens 27, 159–164, doi:10.1097/MNH.0000000000000402 (2018).

27 Balbotkina, E. V. & Kutina, A. V. Structure and Properties of the Glomerular Filtration Barrier in Vertebrates: Role of a Charge in Protein Filtration. Journal of Evolutionary Biochemistry and Physiology 59, 1891–1910, doi:10.1134/S0022093023060017 (2023).

28 Olinger, E., Schwaller, B., Loffing, J., Gailly, P. & Devuyst, O. Parvalbumin: calcium and magnesium buffering in the distal nephron. Nephrol Dial Transplant 27, 3988–3994, doi:10.1093/ndt/gfs457 (2012).

29 Subramanya, A. R. & Ellison, D. H. Distal convoluted tubule. Clin J Am Soc Nephrol 9, 2147–2163, doi:10.2215/CJN.05920613 (2014).

30 Buageaw, A. et al. GDNF family receptor alpha1 phenotype of spermatogonial stem cells in immature mouse testes. Biology of reproduction 73, 1011–1016, doi:10.1095/biolreprod.105.043810 (2005).

31 Grasso, M. et al. Distribution of GFRA1-expressing spermatogonia in adult mouse testis. Reproduction 143, 325–332, doi:10.1530/REP-11-0385 (2012).

32 Garbuzov, A. et al. Purification of GFRalpha1+ and GFRalpha1-Spermatogonial Stem Cells Reveals a Niche-Dependent Mechanism for Fate Determination. Stem Cell Reports 10, 553–567, doi:10.1016/j.stemcr.2017.12.009 (2018).

33 Oulad-Abdelghani, M. et al. Characterization of a premeiotic germ cell-specific cytoplasmic protein encoded by Stra8, a novel retinoic acid-responsive gene. The Journal of cell biology 135, 469–477, doi:10.1083/jcb.135.2.469 (1996).

34 Huang, C. J., Tu, C. T., Hsiao, C. D., Hsieh, F. J. & Tsai, H. J. Germ-line transmission of a myocardium-specific GFP transgene reveals critical regulatory elements in the cardiac myosin light chain 2 promoter of zebrafish. Dev Dyn 228, 30–40, doi:10.1002/dvdy.10356 (2003).

35 Herzlinger, D. & Hurtado, R. Patterning the renal vascular bed. Seminars in cell & developmental biology 36, 50–56, doi:10.1016/j.semcdb.2014.08.002 (2014).

36 Hurtado, R. et al. Pbx1-dependent control of VMC differentiation kinetics underlies gross renal vascular patterning. Development 142, 2653–2664, doi:10.1242/dev.124776 (2015).

